# Predicting gait adaptations due to ankle plantarflexor muscle weakness and contracture using physics-based musculoskeletal simulations

**DOI:** 10.1101/597294

**Authors:** Carmichael F. Ong, Thomas Geijtenbeek, Jennifer L. Hicks, Scott L. Delp

**Affiliations:** Department of Bioengineering, Stanford University, Stanford, California, United States of America; Department of Biomechatronics & Human-Machine Control, Delft University of Technology, Delft, The Netherlands; Department of Mechanical Engineering, Stanford University, Stanford, California, United States of America; Department of Orthopaedic Surgery, Stanford University, Stanford, California, United States of America

## Abstract

Deficits in the ankle plantarflexor muscles, such as weakness and contracture, occur commonly in conditions such as cerebral palsy, stroke, muscular dystrophy, and Charcot-Marie-Tooth disease. While these deficits likely contribute to observed gait pathologies, elucidating cause-effect relationships is difficult due to the often co-occurring biomechanical and neural deficits. To elucidate the effects of weakness and contracture, we systematically introduced isolated deficits into a musculoskeletal model and generated simulations of walking to predict gait adaptations due to these deficits. We developed a planar model containing 9 degrees of freedom and 18 musculotendon actuators, and an optimization framework through which we imposed simple objectives, such as minimizing cost of transport while avoiding falling and injury, and maintaining head stability. We first validated that our model could generate gaits that reproduced experimentally observed kinematic, kinetic, and metabolic trends for two cases: 1) walking at prescribed speeds between 0.50 m/s and 2.00 m/s and 2) walking at self-selected speed. We then applied mild, moderate, and severe levels of muscle weakness or contracture to either the soleus (SOL) or gastrocnemius (GAS) or both of these major plantarflexors (PF) and retrained the model to walk at a self-selected speed. The model was robust to all deficits, finding a stable gait in all cases. Severe PF weakness caused the model to adopt a slower, "heel-walking" gait. Severe contracture of only SOL or both PF yielded similar results: the model adopted a "toe-walking" gait with excessive hip and knee flexion during stance. These results highlight how plantarflexor weakness and contracture may contribute to observed gait patterns. Our model, simulation and optimization framework, and results are freely shared so that others can reproduce and build upon our work.

**Author summary:** Deficits in the muscles that extend the ankle are thought to contribute to abnormal walking patterns in conditions such as cerebral palsy, stroke, muscular dystrophy, and Charcot-Marie-Tooth disease. To study how deficits in these muscles contribute to abnormal walking patterns, we used computer simulations to systematically introduce muscle deficits into a biomechanically accurate model. We first showed that our model could discover realistic walking patterns over a wide range of speeds when we posed a simple objective: walking while consuming a minimum amount of energy per distance, maintaining head stability, and avoiding injury. We then used the model to study the effect of two commonly observed problems: muscle weakness and muscle tightness. We found that severe weakness of the ankle extensors caused the model to adopt a slower, “heel-walking” gait, and severe tightness caused the model to adopt a crouched, “toe-walking” gait. These results highlight how deficits in the ankle extensor muscles may contribute to abnormal walking patterns commonly seen in pathological populations.

## Introduction

The ankle plantarflexor muscles play an important role in human walking, and deficits in these muscles are thought to contribute to gait pathologies. Previous work has shown that the plantarflexor muscles are important for generating forward acceleration during the push-off phase of walking [1,2]. Individuals with cerebral palsy, stroke, muscular dystrophy, or Charcot-Marie-Tooth (CMT) disease often exhibit plantarflexor muscle weakness [3–9] and contracture [3,9–15]. However, these muscular deficits often co-occur with other abnormalities, such as skeletal deformities and neural deficits, making it difficult to assess the independent contributions of these muscular deficits to gait abnormalities.

Previous experimental work has sought to better understand the cause-effect relationship between plantarflexor weakness and gait pathologies by performing a tibial-nerve block in one leg in healthy subjects [16]. Adaptations on the affected side included a reduced ability to transfer weight to the contralateral leading limb during terminal stance and a shortened single-limb stance duration. Although this experiment helps to establish a cause-effect relationship, nerve blocks are invasive and thus difficult to repeat for many muscles. Further, these experiments cannot reveal the effects of contracture, which produces abnormally high passive stiffness of muscles.

Musculoskeletal simulations built from experimental gait data have been used to study gait pathologies. For example, simulations of individuals with cerebral palsy have quantified individual muscle contributions to body weight support and forward propulsion [17], the minimum muscle strength required to walk in a crouch gait [18], and the contributions of contracture and spasticity to increased hamstring resistance [19]. These studies suggest strong links between muscle deficits and the observed gait adaptations; however, since these studies tracked experimental data from patients with a combination of muscular, skeletal, and neural deficits, the independent effects of muscular weakness and contracture on the observed gait adaptations cannot be assessed.

Simulations in which kinematics are generated *de novo* (i.e., without tracking experimental data) can help reveal cause-effect relationships between muscular deficits and gait abnormalities. Researchers have created controllers that could generate various gait patterns, including level walking [20–23] and running [21,23], inclined walking [22,23], loaded walking [22], stair ascent, and turning [23]. More recently, Song and colleagues sought to understand which factors contributed to decreased walking performance in the elderly population [24]. By systematically introducing the neural, muscular, or skeletal deficits seen in this population and training each model to walk, they determined that decreased muscle strength and mass of all muscles contributed most to typically observed gait adaptations. Although simulations of this type are valuable, they are not yet broadly used because it is challenging to generate simulations *de novo* and to reproduce previous work. Overcoming these challenges has been difficult because of the sensitivity of forward simulations to a software environment and limited sharing of software and models. More work is also needed to validate that simulations of this type capture the adaptations observed in experiments (e.g., due to varying walking speeds), before adjusting the model to generate simulations under new conditions that cannot be observed in controlled experiments.

The goal of our work was to determine which gait adaptations arise from weakness or contracture of the plantarflexor muscles. To this end, we first created and validated an optimization framework and musculoskeletal model that could generate realistic motions *de novo*. Our controller followed the previously described reflex-based controllers [20,21,23], and the parameters of this controller were iteratively updated within the optimization framework. We validated our results over a wide range of prescribed walking speeds and at self-selected speed; this step was necessary as low gait speed is commonly observed in individuals with gait impairments. We then introduced weakness or contracture in the plantarflexor muscles, generated new gait patterns, and analyzed subsequent changes in the kinematics and kinetics of gait. Our model and results are freely available at https://simtk.org/projects/pfdeficitsgait along with a Docker build file and setup files so others can reproduce our work. We also provide downloads for the control and optimization framework (SCONE, Simulation and CONtrol Environment) and accompanying documentation at http://scone.software to enable others to use and build upon our simulation framework.

**Fig 1.**
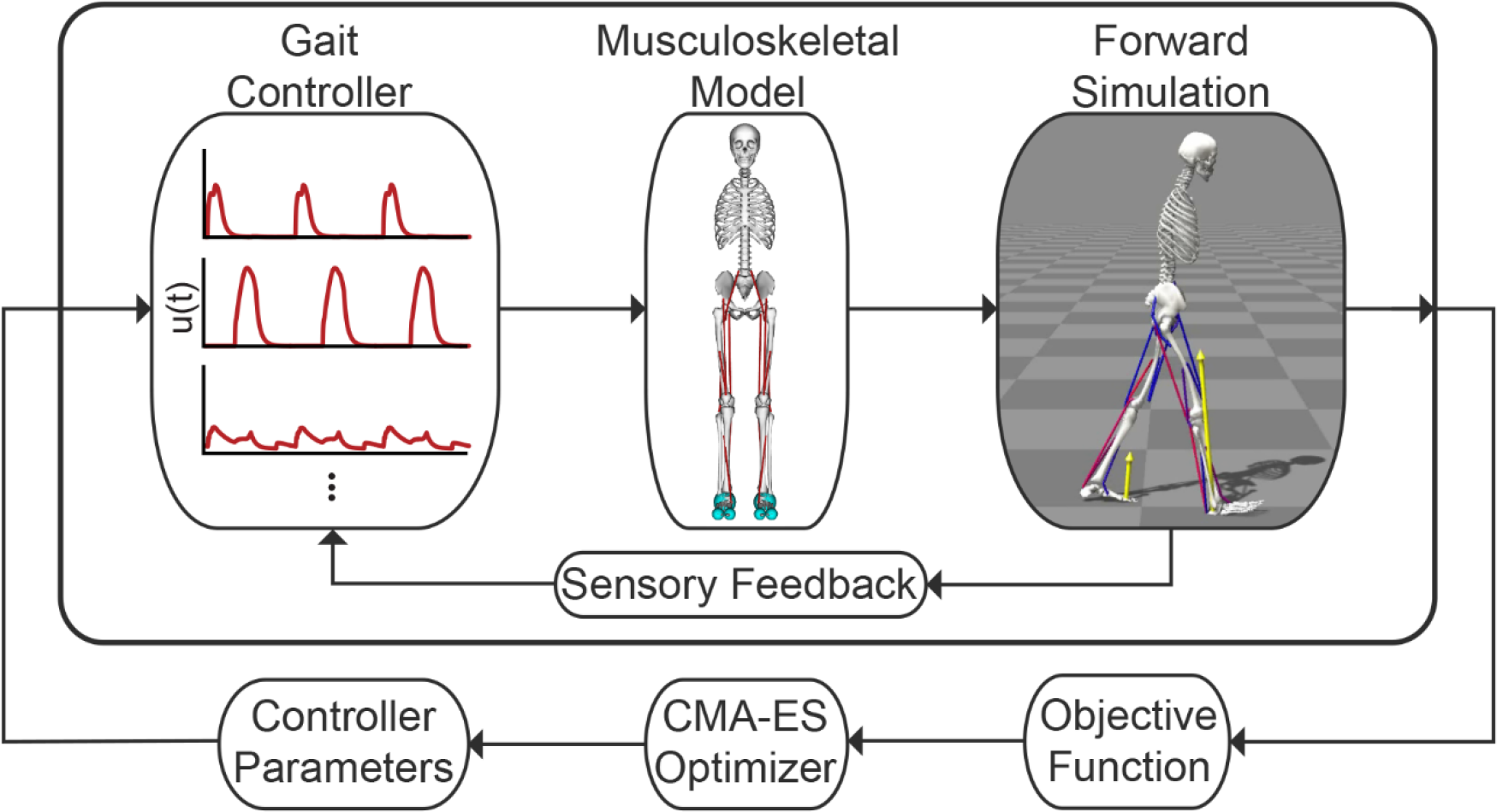
Single shooting framework for dynamic optimization. We trained a planar, musculoskeletal model actuated by 18 Hill-type musculotendon actuators by optimizing the parameters of a gait controller based on an objective function that sought to minimize metabolic cost, avoid falling and injury, and stabilize the head. A gait controller computed muscle excitations, u(t), for a musculoskeletal model to generate a forward simulation. Sensory feedback, based on the model’s muscle and joint states, was used in a feedback loop with the gait controller. The objective function quantified the performance of each simulation, and a Covariance Matrix Adaptation Evolutionary Strategy (CMA-ES) optimization method updated the values of the variables in the optimization problem.

## Results and Discussion

We generated our simulations using a single shooting method (Fig 1), which is discussed in more detail in the Methods section. We first compare our simulations of unimpaired walking to experimental kinematic, kinetic, and metabolic data. We then detail how we modeled plantarflexor weakness and contracture and discuss the kinematic and kinetic adaptations caused by introducing these deficits into our musculoskeletal model. Video highlights of our results are provided in S1 Video, and videos of all simulations discussed here are provided in S2 Video.

### Validating the model’s gait over a range of speeds

We validated the ability of our model and optimization framework to capture trends in kinematics, kinetics, and spatio-temporal measures when walking at different speeds by generating seven simulations of gait at prescribed speeds between 0.50 m/s and 2.00 m/s, at intervals of 0.25 m/s. All simulations found a steady gait pattern at a speed within 0.05 m/s of the speed prescribed by the optimization’s objective function (S2 Video). Individual comparisons of each speed to experimental data [25] are provided in S1 Fig.

**Fig 2.**
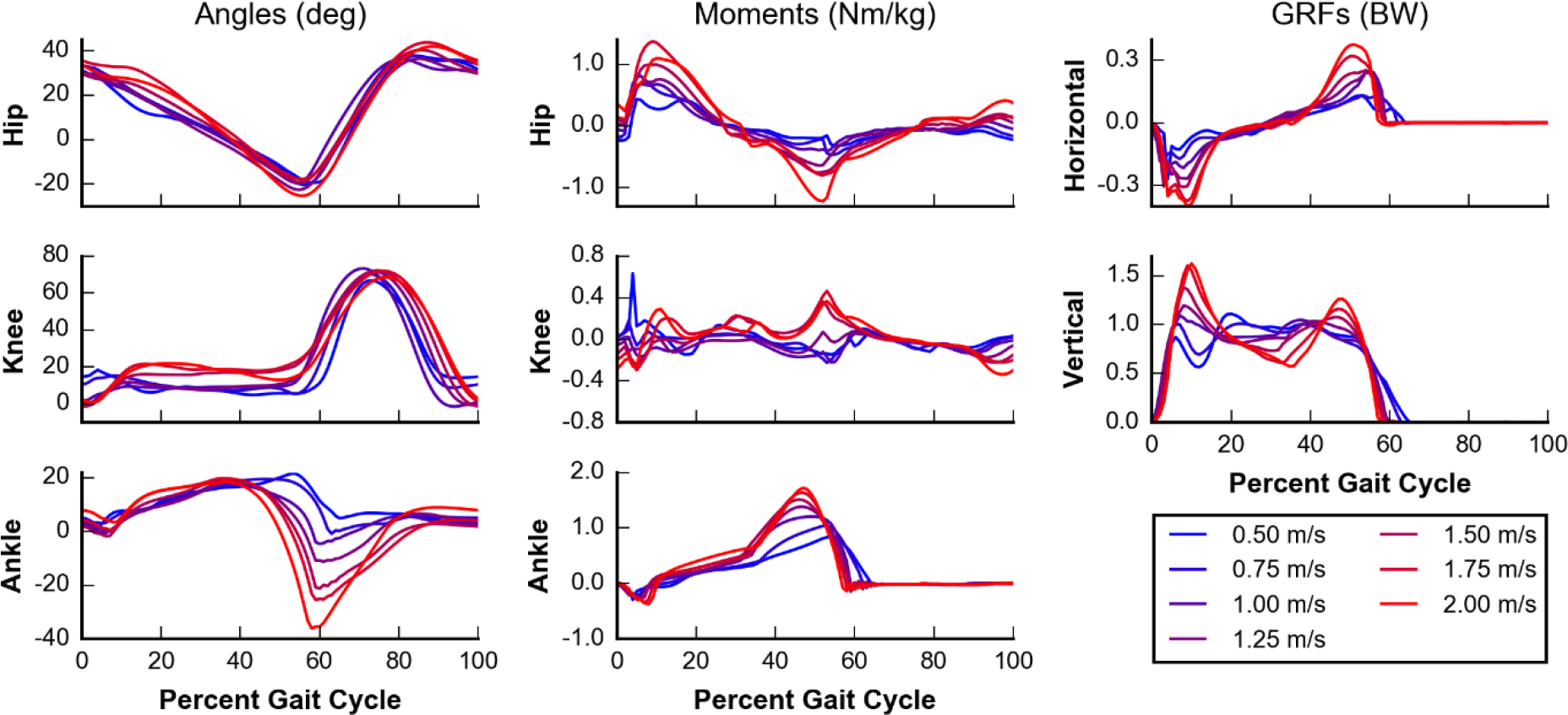
Kinematics and kinetics of simulated walking over a range of speeds. Seven prescribed speeds between 0.50 m/s (blue) and 2.00 m/s (red) at intervals of 0.25 m/s were analyzed. Joint angles (left column) and joint moments (middle column) are plotted for the hip (top row), knee (middle row), and ankle (bottom row). Positive joint angles indicate flexion, and positive joint moments indicate extension. Ground reaction forces (GRFs) (right column) are shown for both horizontal (top row) and vertical (middle row) directions. With exceptions at the knee joint, we observed trends that occur in experimental data [25], including greater joint angle ranges, joint moments, and ground reaction forces at higher speeds.

Simulated kinematic and kinetic adaptations, including joint angles, joint moments, and ground reaction forces (GRFs), matched trends observed in experimental data [25] with increasing speeds (Fig 2). Hip and ankle joint angle excursions increased with increasing speeds. Kinetic data, such as peak flexion and extension moments about the hip, knee, and ankle, and the peak horizontal and vertical GRFs, also increased with increasing speeds. Some experimental trends, however, were not captured in our simulations. The knee joint angle range and peak flexion during swing did not increase at higher speeds. At the slowest prescribed speeds (0.50 m/s and 0.75 m/s), the GRFs show a slow resonance not seen in experimental data. This was likely due to our choice for contact parameters, which yielded a softer contact than previous work [22]. While this choice likely contributed to resonance, it also sped simulations up by roughly 2- to 3-fold. We note one aberrant point for the simulation at the slowest speed (0.50 m/s) in the knee joint moment plot that shows a spike of knee extension in early stance, but it does not appear to affect other kinematic or kinetic measures. Overall, while these results show that our simulations capture many salient kinematic and kinetic trends, we must be cautious in interpreting GRF results at speeds at or below 0.75 m/s.

**Fig 3.**
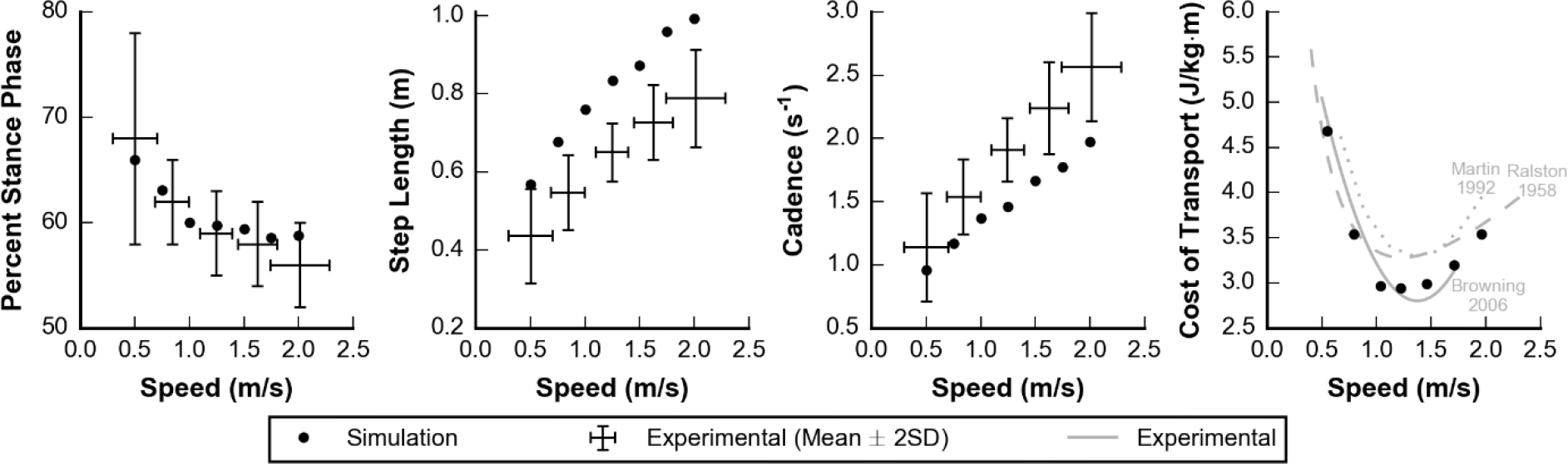
Spatio-temporal and metabolic parameters over a range of speeds. Simulated (dots) and experimental (lines) data are plotted for percent stance phase (left), step length (middle left), cadence (middle right), and cost of transport (right). Experimental data for the left three panels are shown with 2 standard deviations [25]. In the right panel, the magnitude and characteristic shape of the cost of transport bowl were similar between the simulated data and experimental data sets [26–28].

Percent stance phase (i.e., the percent of time each foot is in stance during a gait cycle) decreases, and step length and cadence increased with increasing speeds as observed experimentally [25–28] (Fig 3). Percent stance phase was within 2 standard deviations (SD) of experimental data at all speeds. Although the trends of step length and cadence follow experimental data, the simulations generated gaits with a longer step length and a slower cadence than observed experimentally. Our simulations were also able to reproduce experimentally observed trends in metabolic cost of transport over the range of speeds we tested (Fig 3, right panel). In particular, a greater increase in cost of transport was observed when walking slower than optimal as compared to faster than optimal.

### Validating self-selected gait of the model

We tested the ability of our model and optimization framework to generate a realistic gait at self-selected speed by generating simulations that did not have a prescribed speed within the optimization’s objective function. Instead, we imposed a minimum speed of 0.75 m/s, which helped guide the optimizer to better solutions but was sufficiently low that it did not affect the final optimized solutions. We generated 3 simulations by initializing the optimization framework with the optimized parameters for 3 of the prescribed speeds from the previous section: slowest (0.50 m/s), middle (1.25 m/s), and fastest (2.00 m/s) speeds. The optimization yielded 3 simulations with similar speeds of 1.23 m/s, 1.21 m/s, and 1.29 m/s, respectively. Each speed was within 2 SDs of the expected self-selected speed (1.25 ± 0.15 m/s) for an individual of our model’s leg length [25]. This suggests that our optimization framework can robustly find solutions and is not sensitive to initial guess. For simplicity, only the simulation initialized with the middle speed is described in the remainder of this section.

Our simulated kinematic and kinetic trajectories, including joint angles, joint moments, and GRFs, were similar both in value and shape to experimental data for self-selected gait (Fig 4). For the majority of the gait cycle, simulated trajectories were within 2 SDs of experimental data describing self-selected gait. For each of the trajectories, the root-mean-squared error (RMSE) between simulated and experimental data was no more than 1.55 SD, and for each trajectory except the knee extension moment, the normalized cross-correlation (NCC) [29], which measures shape similarity between the simulated and experimental data sets, was at least 0.82 (Table 1).

The three trajectories with the highest RMSEs, hip joint angle (1.55 SD), knee joint angle (1.48 SD), and knee joint moment (1.46 SD), each contained times when the simulation deviated by more than 2 SDs from the experimental mean. The hip and knee joint angles showed excessive flexion during the swing phase. These were likely compensations due to the simplifications in the model, as degrees of freedom that contributed to foot clearance, such as pelvic list and hip abduction, were not included in the model. Despite the larger RMSEs, the high NCCs of the simulated hip (0.95) and knee (0.97) joint angles to their corresponding experimental trajectories indicate the shape of the simulated and experimental trajectories matched well. The simulated and experimental knee extension moments did not have a similar shape, with a NCC of -0.04. However, there were large standard deviations in the experimental data during this part of the gait cycle, which is not considered when computing NCC. Differences during early stance explain this discrepancy in knee joint moment trajectories, as the model generated a knee flexion moment when experimental data exhibits a knee extension moment. The model took long steps and landed with an extended knee, causing the GRF to generate a knee extension moment, rather than a knee flexion moment in early stance. Thus, the muscles had to generate a knee flexion moment, rather than a knee extension moment, to prevent hyperextension.

**Fig 4.**
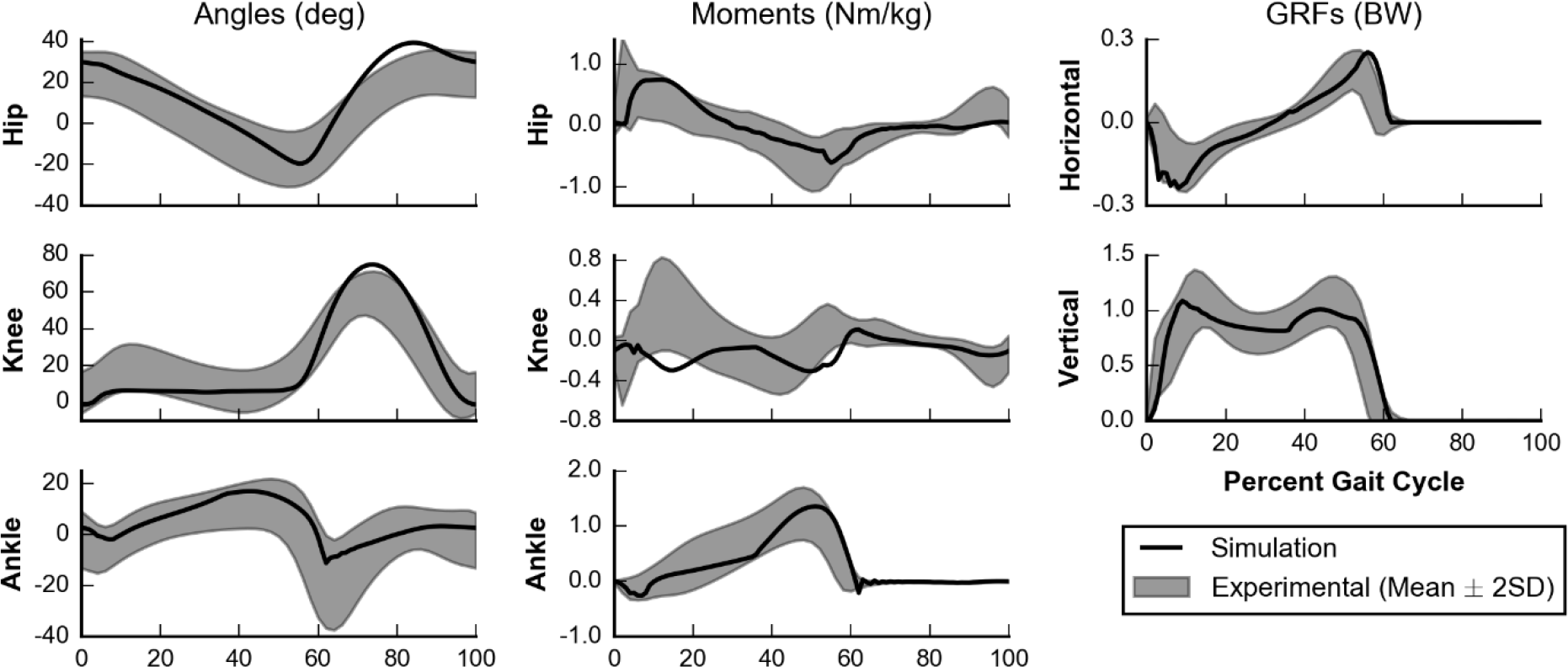
Kinematics and kinetics of self-selected gait. Simulated (black line) kinematics and kinetics are compared to experimental data (gray area) [25]. Joint angles (left column) and joint moments (middle column) are plotted for the hip (top row), knee (middle row), and ankle (bottom row). Ground reaction forces (right column) are shown for both horizontal (top row) and vertical (middle row) directions. Positive angles indicate flexion, and positive moments indicate extension. Note that the experimental data for hip flexion angle was shifted by 11.6° to account for the difference in pelvis orientation definitions between the experimental data set and the musculoskeletal model.

**Table 1.** Similarity metrics between simulated and experimental data for self-selected gait.

|  | Angles |  |  | Moments |  |  | GRFs (stance only) |  |
| --- | --- | --- | --- | --- | --- | --- | --- | --- |
|  | Hip | Knee | Ankle | Hip | Knee | Ankle | Horizontal | Vertical |
| **Root-mean-squared error (SD)** | 1.55 | 1.48 | 0.82 | 1.08 | 1.46 | 1.17 | 1.04 | 0.78 |
| **Normalized cross-correlation** | 0.95 | 0.97 | 0.82 | 0.83 | -0.04 | 0.95 | 0.95 | 0.99 |

Root-mean-squared error compares simulation mean trajectories to experimental mean and standard deviation, reported in units of standard deviation (SD). Given the experimental mean and standard deviation over the gait cycle, we compute a Z-score for the simulation data at every 1% of the gait cycle. Then, we take the square root of the sum of squares of all the Z-scores. Normalized cross-correlation compares mean trajectories between simulation and experimental data [25].

### Simulating walking with plantarflexor weakness or contracture

The results of the previous two sections gave us confidence that this model and optimization framework could generate realistic gaits over a wide range of speeds, and could self-select a realistic gait and speed. We then used the framework to study how gait would adapt to deficits in the plantarflexors by adjusting the model’s parameters and using the same optimization framework.

We modeled mild, moderate, and severe weakness by reducing a muscle’s maximum isometric force 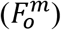 to 25%, 12.5%, and 6.25%, respectively, of its original value. The resulting values for 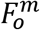 were similar to those used in a previous study that computed the minimum plantarflexor muscle strength needed to walk either normally or in a crouch [18]. We modeled mild, moderate, and severe contracture by reducing a muscle’s optimal fiber length 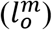 to 85%, 70%, and 55%, respectively, of its original value. The severe case was based on experimental data [30] and previously used as a model of contracture [31]. We separately applied each of these 6 deficits to the soleus only (SOL), gastrocnemius only (GAS), or both plantarflexors (PF), to yield 18 total deficit cases.

**Fig 5.**
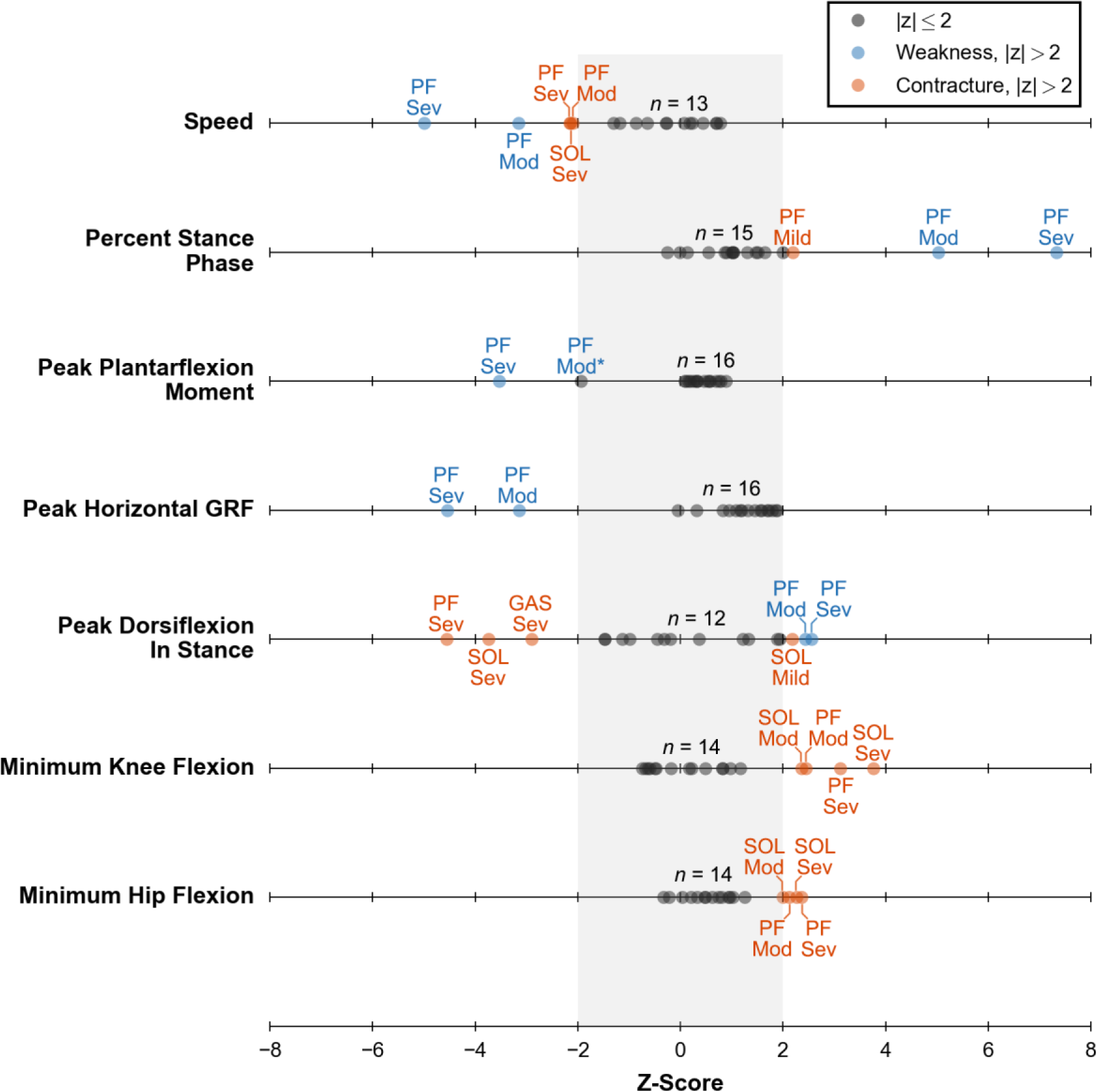
Z-scores of key kinematic and kinetic measures of gait for all deficit cases. Z-scores were computed using previous experimental data of individuals walking at a self-selected speed [25]. All 18 cases (dots) are plotted for each measure. A black dot indicates any case within 2 SDs of normal (gray band). A blue or orange dot indicates either a weakness or contracture case, respectively, which is outside of 2 SDs. All of these cases are labeled with the affected muscle or muscles: soleus (SOL), gastrocnemius (GAS), or both (PF); and severity level: mild (Mild), moderate (Mod), or severe (Sev). One other case is labeled (see Peak Plantarflexion Moment, PF Mod*) as it was a clear outlier compared to other deficit cases. The count above each line (e.g., *n* = 13) indicates the number of unlabeled dots.

The optimization framework generated a stable gait in all deficit cases, highlighting its ability to change neural control in response to weakness and contracture (S2 Video). Furthermore, the model’s gait was robust to most deficits, and many cases had kinematic and kinetic measures within 2 SDs of walking at a self-selected speed (Fig 5, S2 Fig). By count, the most affected cases were moderate PF weakness, severe PF weakness, severe SOL contracture, and severe PF contracture. In the following sections, we detail the weakness and contracture cases that caused the model to adopt a gait with kinematic or kinetic features outside of 2 SDs of normal walking at a self-selected speed. We provide complete kinematic and kinetic trajectories for all simulations of muscle weakness and contracture in S3 Fig.

### Walking with plantarflexor weakness

Plantarflexor weakness decreases the capacity of muscles to generate ankle plantarflexion moments, which are critical for generating forward propulsion during walking [1,2,16]. Our simulations supported this notion as plantarflexion moments during walking were most affected by moderate and severe PF weakness and minimally affected by SOL or GAS weakness only (Fig 5, third row). This suggests that in cases of weakness to only the SOL or GAS, the unimpaired plantarflexor had sufficient capacity to compensate for weakness in the other muscle. For moderate and severe PF weakness, peak ankle plantarflexion moment decreased by 43% and 69%, respectively, when compared to simulated unimpaired walking (Fig 6, bottom row). This highlights the reduced moment generating capacity in these cases.

Other gait parameters, such as peak horizontal GRF, self-selected walking speed, and percent stance were abnormal for moderate and severe PF weakness as well (Fig 5). Compared to simulated unimpaired walking, peak horizontal GRF decreased by 55% and 70% for moderate and severe PF, respectively, which led to slower walking speeds of 1.01 m/s and 0.87 m/s, respectively. These decreases in gait speed in response to weakness are unsurprising given the direct relationship between the push-off phase and walking speed. Reduced walking speed is commonly observed in individuals with cerebral palsy [32], stroke [33], muscular dystrophy [34], and CMT disease [35], where plantarflexor weakness is common. After accounting for these reduced walking speeds, the model spent an increased percentage of the gait cycle in stance phase, 69% and 74% with moderate and severe PF weakness, respectively. For comparison, simulated unimpaired walking at a slower speed of 0.75 m/s had a percent stance phase of only 63% (Fig 3). These large increases in percent stance phase are consistent with trends seen in the pathological population. For example, CMT patients who had a “plantar flexor strength deficit” were found to adopt a slower gait with an increased time spent in stance phase compared to the gait chosen by patients without the strength deficit [36].

**Fig 6.**
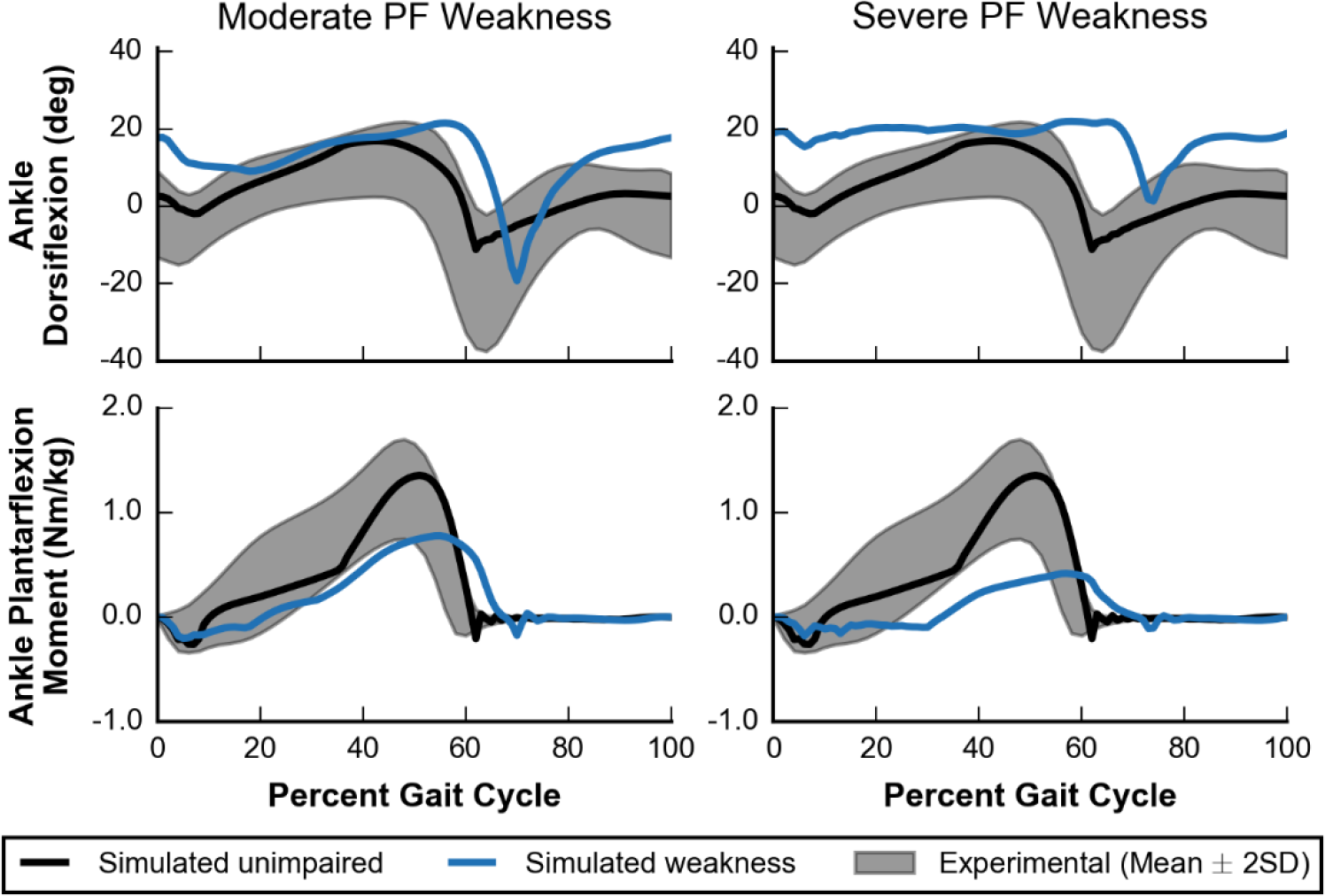
Ankle kinematics and kinetics for moderate and severe PF weakness. Simulated ankle kinematics and kinetics for unimpaired walking (black lines) and walking with PF weakness (blue lines) are compared to experimental data (gray area) [25]. Ankle dorsiflexion (top) and ankle plantarflexion moment (bottom) are plotted for cases of moderate (left) and severe (right) PF weakness.

Dramatic kinematic compensations at the ankle were observed at the ankle in cases of moderate and severe PF weakness. These models adopted a calcaneal, or “heel-walking”, gait, landing with substantially increased ankle dorsiflexion that was maintained throughout stance (Fig 6, top row). When compared with simulated unimpaired walking, the model with moderate or severe PF weakness had an increased ankle dorsiflexion at initial contact of 15° and 16°, respectively, and an increased average ankle dorsiflexion during stance of 7° and 11°, respectively. This type of gait is often observed in patients who have received either an intramuscular aponeurotic recession of the plantarflexors or Achilles tendon lengthening to correct contracture [37–39], and it is commonly thought that overlengthening during these procedures can lead to plantarflexor weakness [40,41]. Our simulations support the notion that plantarflexor weakness alone could cause calcaneal gait.

While it has been previously hypothesized that plantarflexor weakness may contribute to individuals adopting a crouch gait [18,38,39,41], our simulations did not support this hypothesis. Crouch gait is characterized by increased knee flexion during stance and commonly occurs with increased hip flexion during stance, but our optimization framework found gaits that did not require substantial plantarflexion moment and did not have increased knee and hip flexion during stance (Fig 5, bottom two rows). Our results also differ from previous work that observed a mild crouch with weakened plantarflexors [21], but a key difference was that our simulations did not have a constrained speed whereas the previous work had a prescribed speed of 1.25 m/s. Although our work does not support the notion that plantarflexor weakness alone will lead to crouch gait, it does not rule out that it could be a contributing factor. For example, it might be that plantarflexor weakness is often associated with weakness and contracture in other muscle groups, and a combination of these deficits along with impaired neural control may lead to crouch gait.

### Walking with plantarflexor contracture

Plantarflexor contracture increases passive ankle plantarflexion moments, shifting the ankle’s resting position to a more plantarflexed state and limiting ankle range of motion [11,13]. All cases of severe contracture (i.e., SOL, GAS, and PF) had significantly decreased peak values of dorsiflexion in stance (Fig 5, fifth row). One case, mild SOL contracture, appeared to have an opposite effect, but we believe this is an anomaly as the overall ankle position trajectory did not differ much from the unimpaired case (S3D Fig). Models with severe contracture landed on their toes as the ankle was plantarflexed beyond 0° at initial contact, and stayed on their toes throughout stance (Fig 7, top row). Equinus, or “toe-walking,” gait is commonly observed in individuals with cerebral palsy [42,43]. Plantarflexor contracture is thought to be a major contributing factor [11,44], and our results support this theory.

**Fig 7.**
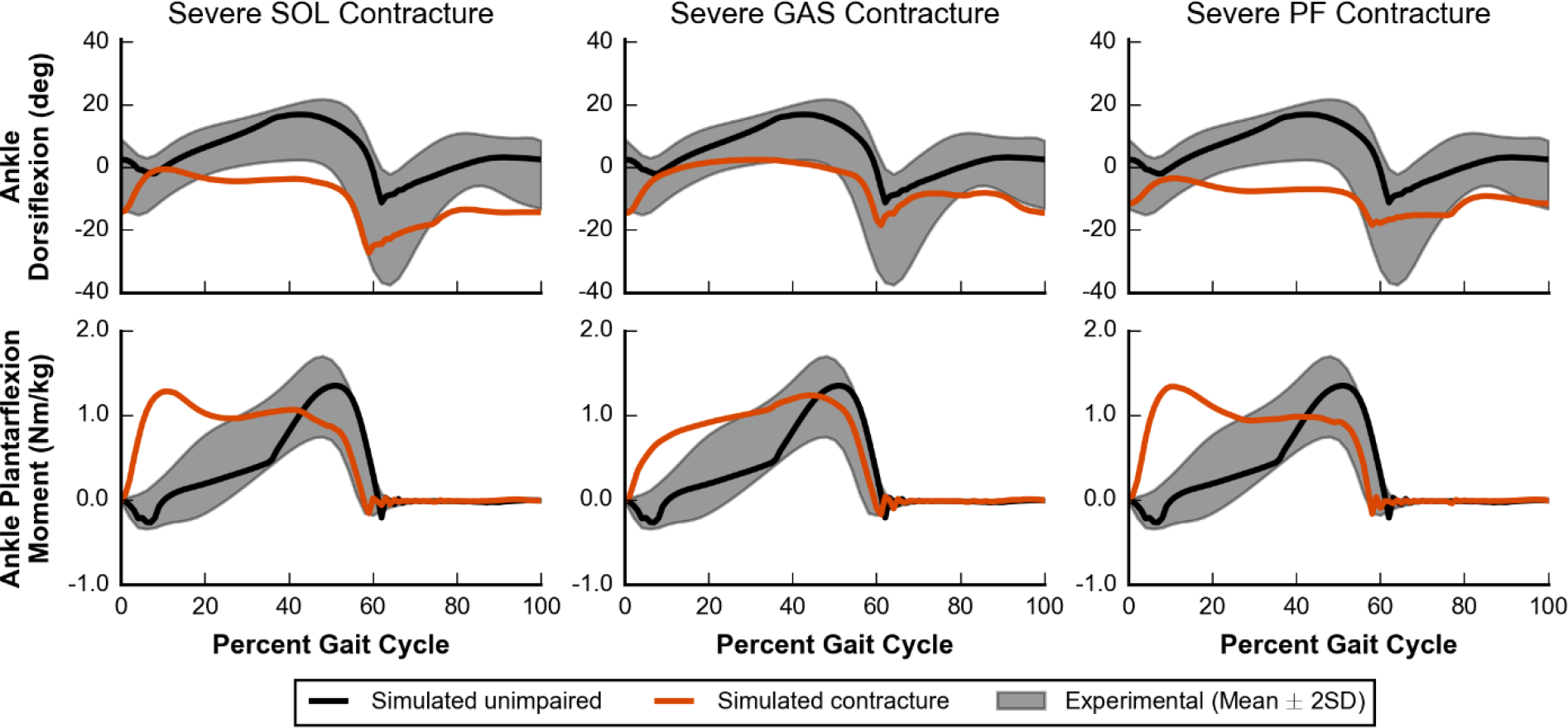
Ankle kinematics and kinetics for severe contracture. Simulated ankle kinematics and kinetics for unimpaired walking (black lines) and walking with severe contracture (orange lines) are compared to experimental data (gray area) [25]. Ankle dorsiflexion (top) and ankle plantarflexion moment (bottom) are plotted for cases of SOL (left), GAS (middle), and PF (right) contracture.

All cases of severe contracture had significantly increased plantarflexion moments during early stance (Fig 7, bottom row), which was expected since an equinus gait necessitates large plantarflexion moments at initial contact to prevent excessive dorsiflexion. In particular, severe SOL or PF contracture had a peak plantarflexion moment during early stance rather than during the push-off phase. Although peak values were largely unaffected, plantarflexion moments during the push-off phase did decrease which led to a subsequent decrease in peak horizontal GRF; all contracture cases had a peak horizontal GRF that was between 2% and 23% lower than our simulated unimpaired case (Fig S2, fourth row).

**Fig 8.**
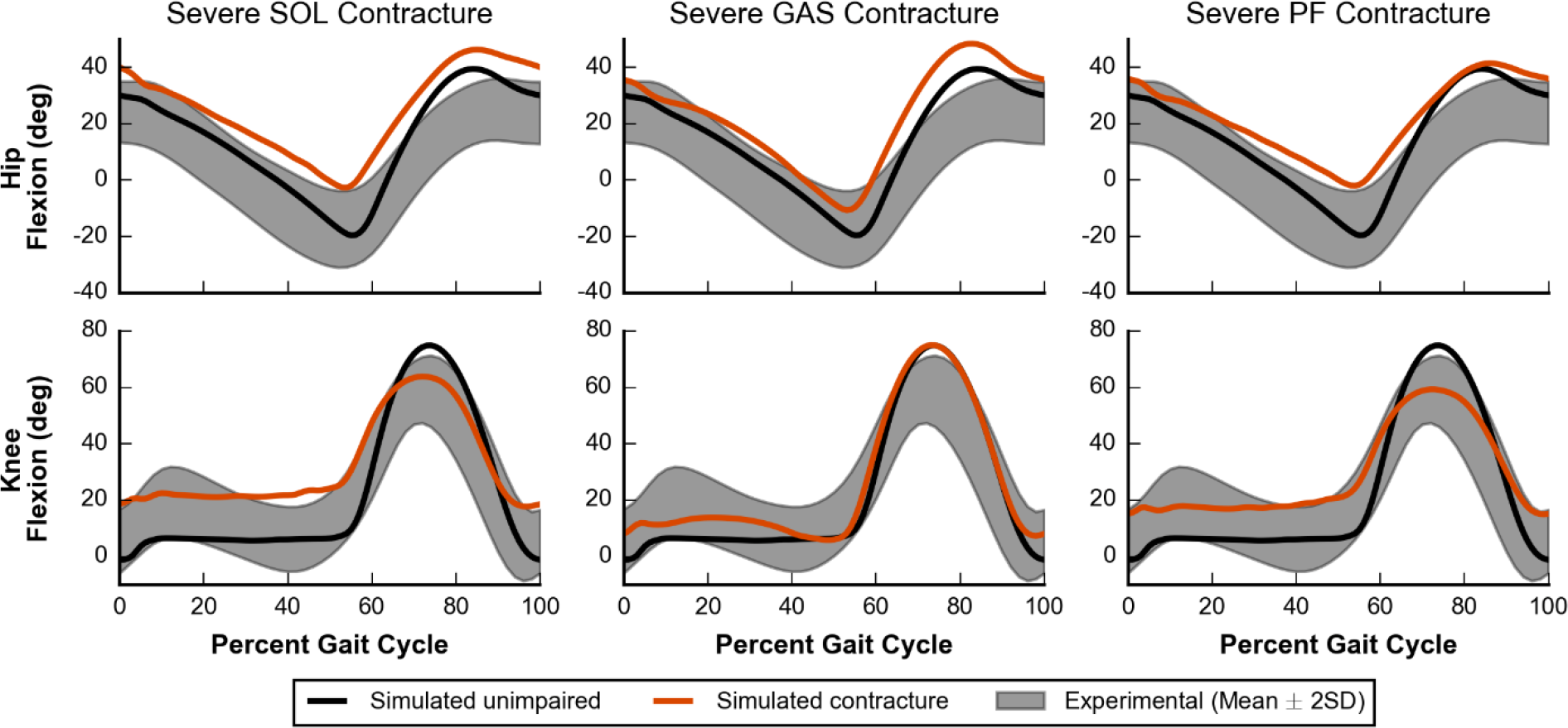
Hip and knee kinematics for severe contracture. Simulated hip and knee kinematics for unimpaired walking (black lines) and walking with severe contracture (orange lines) are compared to experimental data (gray area) [25]. Hip flexion (top) and knee flexion (bottom) are plotted for cases of SOL (left), GAS (middle), and PF (right) contracture.

As severity of contracture increased, models adapted by walking slower and with a shorter stance phase. The three slowest conditions, which included severe SOL, moderate PF, and severe PF contracture, all walked at a significantly slower speed of 1.08 m/s with a slightly smaller percent stance phase of 59% (Fig 5, top two rows). This small decrease in percent stance phase agrees well with experimental data of individuals walking in equinus [45].

In cases of severe SOL, GAS, and PF contracture, the model adopted gaits that were more crouched throughout the whole stance phase, as measured by increased knee and hip flexion, when compared to simulated unimpaired walking (Fig 8). Cases of SOL and PF contracture were more crouched than cases of GAS contracture (S2 Fig, bottom two rows). These results highlight two conclusions. First, contracture in the plantarflexor muscles can lead to dramatic compensations at joints that they do not span. Second, because contracture of only GAS had the smallest effect, our results suggest that the crouched postures adopted by the model were affected less by the GAS directly flexing the knee and more by walking in equinus.

### Limitations and future work

Simplifications in the model may have affected the observed kinematic trajectories. The model was constrained to sagittal plane motion and consequently lacked degrees of freedom, such as hip abduction, pelvis list, and pelvis rotation, which allow for out-of-plane motions. This made foot clearance more difficult and was likely the cause of compensations in the sagittal plane that increased knee and hip flexion during swing. For the models with deficits, these planar constraints did not allow for commonly observed compensations that are out of the sagittal plane, such as hip circumduction [46,47].

Although we chose to model weakness by decreasing 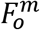 and contracture by decreasing 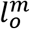, there are other ways to model these deficits. For instance, weakness could also be modeled as decreased maximum muscle excitation, which may be important in cases such as stroke [48], and contracture could also be modeled as decreased tendon slack length [49] or increased passive muscle stiffness [50]. These alternate modeling approaches may have led to different gait compensations, and future work to test sensitivity of our results to variations in modeling choices would be valuable.

Although our objective function contained terms that are thought to capture high-level goals of walking, such as minimizing cost of transport [51] while maintaining head stability [52] (see Methods), it may not comprehensively represent goals of human walking, especially amongst individuals with gait pathologies. For example, some individuals may choose a trade-off between metabolic efficiency and joint loading or stability to perturbations. Furthermore, the relative trade-off between these competing objectives likely varies between individuals.

Due to the non-convex nature of the optimization problem and the stochastic nature of our optimizer, the solutions provided here are likely to be local minima rather than global minima. To gain confidence in our solutions, we followed previous work that used multiple optimizations from the same seed and restarted optimizations from the previous best result [23,53], and furthermore, we demonstrated how our solution was robust to different initial seeds (see section on “Validating self-selected gait of the model”). This strategy, however, was computationally expensive, and further work is needed to better search this complex, high-dimensional space. Since a good starting guess can be important for finding reasonable results, we provide our results to assist other researchers in future studies.

Although this work presents a validated framework for studying how isolated changes in muscle parameters may lead to changes in gait, future work on how gait adapts with changes in neural control will further our understanding of pathologic gait. Reflex-based controllers, like the one used here, were useful to ensure that the optimization problem was tractable, but a controller like this cannot predict changes in the underlying control structure. Recent advances in the field of reinforcement learning may be able to generate robust simulations with control patterns that do not have a specified and simplified control structure [54,55].

## Conclusion

We present a model and optimization framework suitable for studying sagittal plane adaptations during walking due to muscular deficits. We first validated that, over a wide range of speeds, our model and framework captured many salient features of gait. This framework was then used to study how the model would adapt its gait when plantarflexor weakness or contracture was introduced. Severe plantarflexor weakness caused the model to adopt a calcaneal gait without crouching, while severe plantarflexor contracture caused the model to adopt an equinus gait. Our simulations also showed that contracture of only SOL or both PF caused the model to walk in a more crouched posture than if contracture was applied to only GAS, which suggests that walking in equinus is the major contributor to crouch gait rather than force from the GAS directly flexing the knee. We provide our results freely at https://simtk.org/projects/pfdeficitsgait and software at http://scone.software so that others can easily extend this work.

## Methods

Our optimization framework used a single shooting method to solve the dynamic optimization problem of generating a simulation of gait (Fig 1). We implemented our model in OpenSim 3.3 [56,57] and used an optimization and control framework (SCONE) to implement the gait controller, perform the simulation using OpenSim as the plant, and optimize the parameters of our problem.

### Musculoskeletal model

The musculoskeletal model (Fig 9) was a planar model based on a previous model used to study the lower limb [58]. The musculoskeletal model had 9 degrees of freedom (dof): a 3-dof planar joint connected the pelvis to ground; a 1-dof pin joint represented each hip and ankle; and a 1-dof joint with coupled rotation and translation represented each knee. The lumbar joint was locked at 5° of flexion based on previous walking data [2].

**Fig 9.**
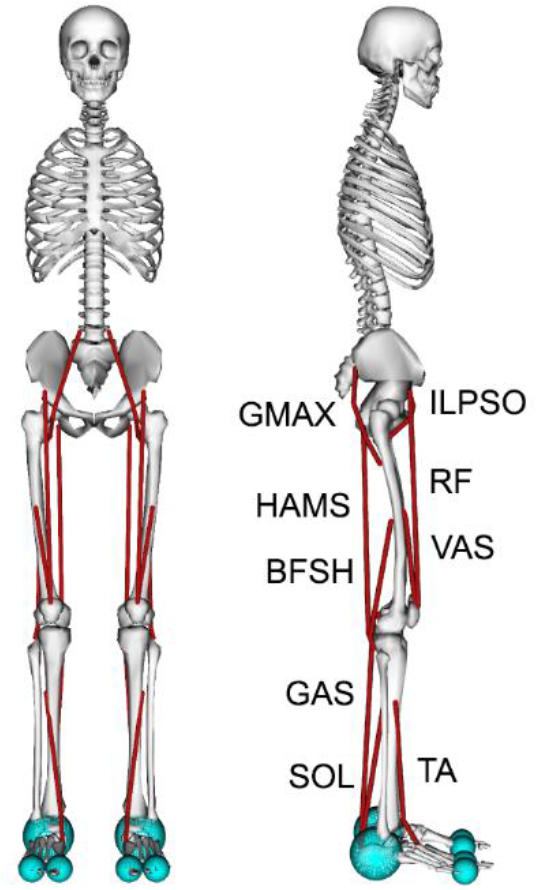
A planar musculoskeletal model for walking. The musculoskeletal model had 9 degrees of freedom (dof): the pelvis-to-ground joint was a 3-dof planar joint, each hip and ankle was a 1-dof pin joint, and each knee was a 1-dof coupled joint. The model’s 18 musclulotendon actuators (red lines) represented the 9 major uniarticular and biarticular muscle groups per leg that drive sagittal plane motion: iliopsoas (ILPSO), gluteus maximus (GMAX), rectus femoris (RF), biarticular hamstrings (HAMS), vasti (VAS), biceps femoris short head (BFSH), gastrocnemius (GAS), tibialis anterior (TA), and soleus (SOL). A compliant contact model was used to generate forces between the spheres at the heel and toes of the feet and the ground plane.

The model was actuated using eighteen Hill-type muscle-tendon units [59], nine per leg. Muscles with similar functions in the sagittal plane were combined into single muscle-tendon units with combined peak isometric forces to represent nine major muscle groups per leg: iliopsoas (ILPSO), gluteus maximus (GMAX), rectus femoris (RF), biarticular hamstrings (HAMS), vasti (VAS), biceps femoris short head (BFSH), gastrocnemius (GAS), tibialis anterior (TA), and soleus (SOL). Peak isometric forces were based on a previous musculoskeletal model [60], whose muscle volumes were based on young, healthy subjects [61]. The tendon strain at peak isometric force was 4.9% [60,62] for all muscles except for the plantarflexors, whose values were set to 10% [63]. To better match passive muscle forces during walking [64], we adjusted the passive force-length curves of the HAMS, VAS, and RF to decrease their passive forces (S1 Table). The tendon slack length of each muscle was calculated based on experimental data [65], as done previously in other musculoskeletal models [60,66]. We set maximum muscle fiber contraction velocity to 15 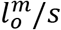 [63,67]. All muscle-tendon parameters are summarized in S1 Table.

Ligaments were modeled as nonlinear, rotational springs that generated torques when a joint was hyperflexed or hyperextended. In our model, ligaments generated torques when the hip was flexed beyond 120° or extended beyond 30°; the knee was flexed beyond 140° or extended beyond 0°; and the ankle was dorsiflexed beyond 20° or plantarflexed beyond 40°.

A compliant contact model [68,69] was used to generate forces between the feet and the ground. Each foot had three contact spheres: one sphere with a radius of 5 cm represented the heel, and two spheres, each with a radius of 2.5 cm, represented the toes. All contact spheres had the following parameters: stiffness coefficient of 500,000 N/m^3/2^, dissipation coefficient of 1.0 s/m, static and dynamic friction coefficients of 0.8, viscous friction coefficient of 0, and transition velocity of 0.1 m/s.

As described in the Results and Discussion, we introduced two types of deficits into the musculoskeletal model: weakness and contracture. We modeled mild, moderate, and severe weakness by reducing the peak isometric force of a muscle 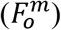 by 25%, 12.5%, and 6.25%, respectively, of the value in the unimpaired model. We modeled mild, moderate, and severe contracture by reducing the optimal fiber length of a muscle 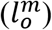 to 85%, 70%, and 55%, respectively, of the value in the unimpaired model. All 6 of these muscle deficits were applied to the SOL, the GAS, or both the SOL and GAS, for a total of 18 cases.

### Gait controller

The gait controller was based on previous reflex-based controllers for human locomotion [20,21,23] and used a combination of a high-level state machine and low-level control laws to calculate muscle excitations. Our controller had 5 high-level states that represented key points in the gait cycle for each leg: early stance (ES), mid-stance (MS), pre-swing (PS), swing (S), and landing preparation (LP). Of the 5 transitions between these high-level states, transitions between 4 of the states were controlled by 4 thresholds that were free parameters in the optimization: 1) ES to MS: horizontal distance between the ipsilateral foot and pelvis was lower than a threshold; 2) PS to S: GRF on the ipsilateral foot was lower than a threshold; 3) S to LP: horizontal distance between the ipsilateral foot and pelvis was greater than a threshold; and 4) LP to ES: GRF on ipsilateral foot was greater than a threshold. The fifth transition, MS to PS, was not controlled by a free parameter and occurred when the contralateral leg entered ES state.

Low-level control laws were active or inactive depending on the high-level controller (Fig 10). There were 5 types of control laws: constant (C), length feedback (L), velocity feedback (V), force feedback (F), and proportional-derivative (PD). All muscle-based feedback laws were positive feedback (i.e., L+, V+, F+) onto the same muscle, except for a negative force feedback (F-) from the soleus to the tibialis anterior. PD control was only used to control the pelvis tilt angle (*θ*) and velocity 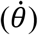 with respect to ground using the muscles crossing the hip joint (i.e., ILPSO, GMAX, and HAMS). Excitations (*u*) are calculated over time (*t*) by the following equations:

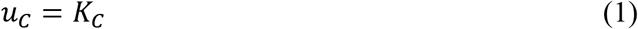

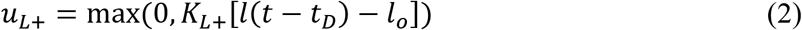

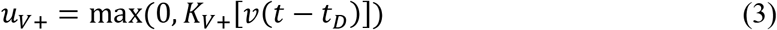

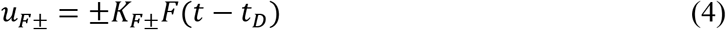

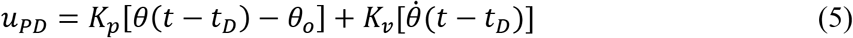

States calculated from the model were muscle length (*l*), muscle velocity (*v*), muscle force (*F*), and pelvis tilt orientation (*θ*) and velocity 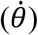. In total, there were 70 free parameters of the optimization, including the controller gains (*K*_*C*_, *K*_*L*+_, *K*_*V*+_, *K*_*F*±_, *K*_*p*_, and *K*_*v*_), and offsets for muscle length feedback (*l*_*o*_) and proportional feedback of *θ* (*θ*_*o*_). For all positive feedback and PD control laws, the parameter for time delay, *t*_*D*_, was set for each muscle depending on the most proximal joint over which the muscle crosses: *t*_*D*_ was 5 ms for the hip, 10 ms for the knee, and 20 ms for the ankle [20]. For the F- law from the soleus to the tibialis anterior, *t*_*D*_ was 40 ms. When considered with a first-order activation time constant of 10 ms, this delay better represented the short-latency TA suppression (56 to 74 ms) observed in experiments [70]. Overall, however, the delays used were still shorter than those measured experimentally [71].

**Fig 10.**
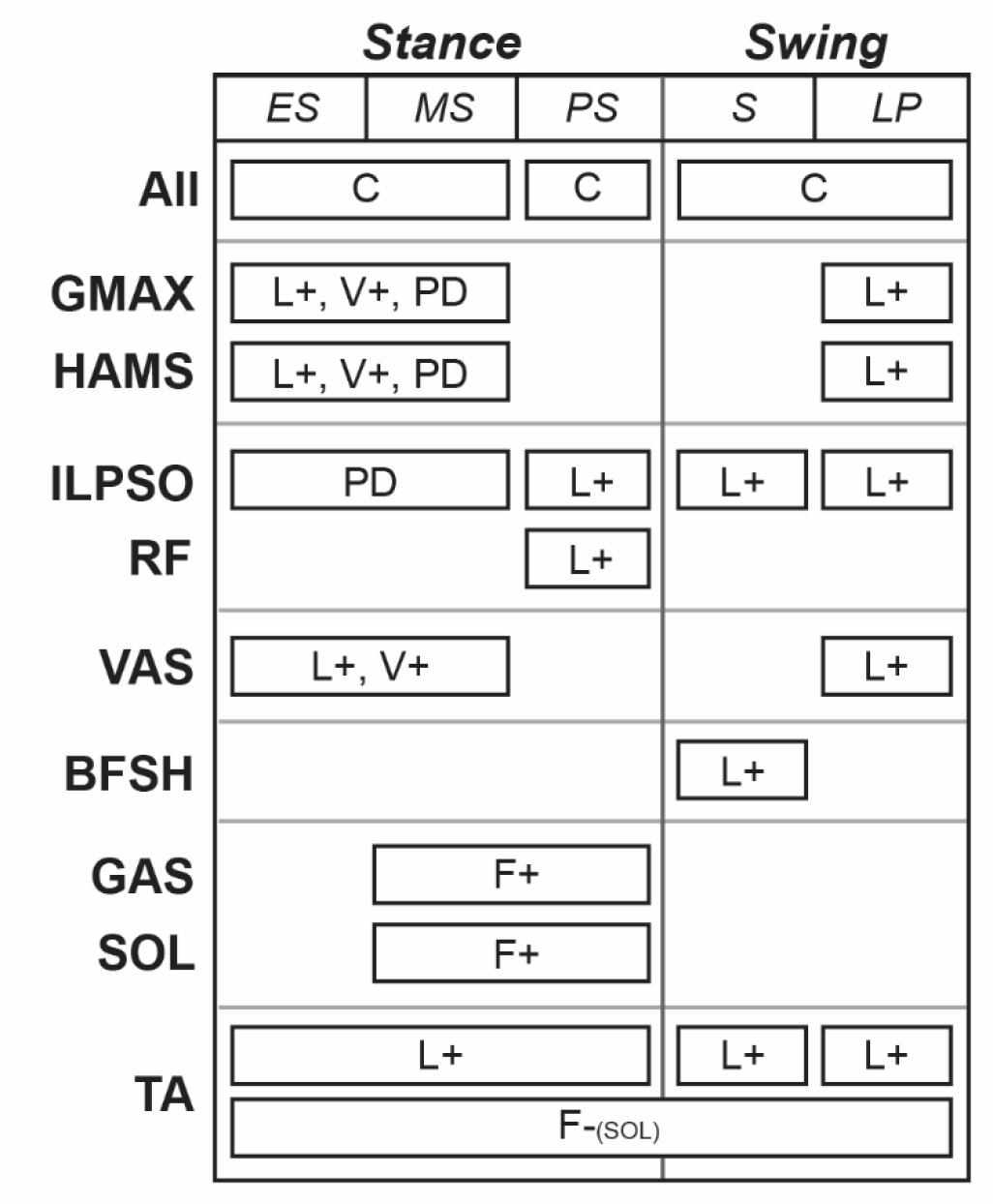
The gait controller used a combination of a state machine and low-level control laws to determine excitations. The state machine had three states in stance (i.e., early stance (ES), mid-stance (MS), and pre-swing (PS)) and two states in swing (i.e., swing (S) and landing preparation (LP)), and it determined when low-level control laws were active. Low-level control laws included constant signals (C) and feedback terms based on muscle length (L), muscle velocity (V), muscle force (F), and proportional-derivative control (PD) based on the pelvis tilt angle. Positive and negative feedback are denoted by (+) and (−), respectively. All feedback laws based on muscle states acted upon the same muscle, except for a negative force feedback from the soleus to the tibialis anterior.

### Simulation and optimization framework

OpenSim was used to form the model’s equations of motion and integrate them [56,57]. The state of the model was initialized by first setting the pelvis’s horizontal position to 0 m. Then, we set initial angular positions and velocities, and the speed of the pelvis according to free parameters in the optimization. Although optimizing velocities may add extra energy into the system, we do not think this affected our results as we found step-to-step variability to be small, and we only analyzed kinematics and kinetics after the first two steps taken. Finally, the vertical position of the pelvis was set such that the GRF was equal to half of the model’s body weight. Integration continued until the desired simulation time was reached or until the model fell, which we defined as the model’s center of mass (COM) height falling below 0.8 times the height of the COM at the start of the simulation.

To evaluate a set of design parameters, we defined an objective function, *J*, which we sought to minimize. The objective, *J,* quantified high-level tasks of walking:

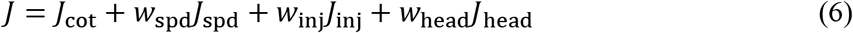

The goal was to minimize the gross cost of transport (*J*_cot_) while maintaining a prescribed step-to-step speed within a range, avoiding falling (*J*_spd_), avoiding ligament injury (*J*_inj_), and stabilizing the head (*J*_head_). To balance competing objectives, the weights were manually adjusted to the following: *w*_spd_ = 10,000, *w*_inj_ = 0.1, and *w*_head_ = 0.25. These weights emphasized finding a gait that did not fall over one that minimized cost of transport, injury, and accelerations at the head. For optimized solutions, the contribution from the *J*_head_ term was larger than those from the *J*_spd_ and *J*_inj_ terms, and thus our results were most sensitive to our choice of *w*head. Because of this, our choice for *w*_head_ represents a trade-off between head stability, especially during early stance, and cost of transport.

*J*_cot_ penalized simulations in which the model walked in a less metabolically efficient manner [51] and was calculated by summing the basal and per-muscle metabolic rates [72,73] to estimate the gross metabolic rate (*Ė*) over the duration of the simulation (*t*_end_), and normalizing by the mass of the model (*m*) and distance travelled (*d*):

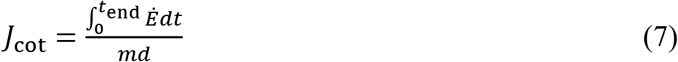

*J*_spd_ penalized simulations in which the model fell at a time (*t*_fall_) before the desired simulation end time (*t*_des_) and had step speeds outside of a prescribed range. *J*_spd_ was calculated using the following equations:

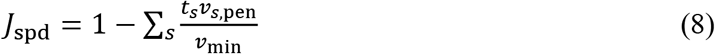

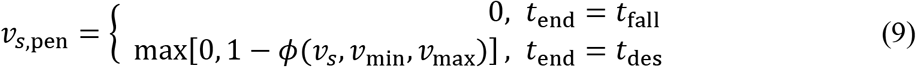

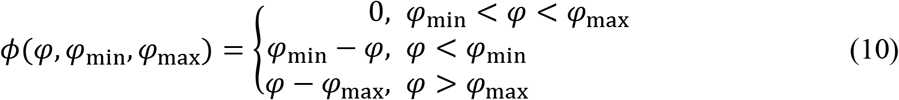

where *s* denotes a step, *t*_*s*_ is the duration of a step, *v*_*s*,pen_ is a penalty for a given step (ranging between 0 and 1, where 1 is no penalty), *v*_*s*_ is the step speed, *v*_min_ and *v*_*max*_ define the minimum and maximum speeds allowed without a penalty, and *Φ* applies a linear penalty if *v*_*s*_ is not between *v*_min_ and *v*_max_. If all steps were within the speed range, *J*_spd_ was 0, and if the model immediately fell, *J*_spd_ was 1. For any given step in which the speed was outside of the range, the step was linearly penalized more heavily (i.e., *v*_*s*,pen_ was closer to 0) the further the speed was out of the range, and its final contribution to *J*_spd_ was scaled by *t*_s_. For simulations with a prescribed speed, the speed range was ±0.05 m/s of the prescribed speed. For example, to prescribe a speed of 1.25 m/s, *v*_min_ = 1.20 m/s and *v*_max_ = 1.30 m/s. For simulations without a prescribed speed, *v*_min_ was 0.75 m/s, and *v*_max_ was arbitrarily large.

*J*_inj_ penalized simulations in which the model engaged the ligaments and was calculated by integrating the sum of joint torques (*T*_*j*_) squared generated by the model’s ligaments:

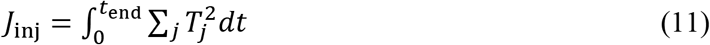

Since head stability is commonly regarded as an important task during gait [74], *J*_head_ penalized simulations in which the model had excessively large accelerations and was calculated by the following equation and Eq 10:

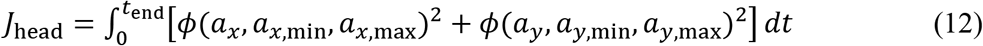

where *a*_*x*_ and *a*_*y*_ are the horizontal and vertical accelerations at a point located near the center of the head, and *a*_*x*,min_, *a*_*x*,max_, *a*_*y*,min_, and *a*_*y*,max_ determine the bounds between which no penalty is applied if *a*_*x*_ and *a*_*y*_ are within the bounds. We set our acceleration bounds as *a*_*x*,min_ = −0.25*g*, *a*_*x*,max_ = 0.25*g*, *a*_*y*,min_ = −0.50*g*, and *a*_*y*,max_ = 0.50*g*, which we approximated from experimental data of human walking [52].

To solve our optimization problem, we used an evolutionary strategy called Covariance Matrix Adaptation Evolutionary Strategy (CMA-ES) with a rank-μ update [75]. In total, there were 90 design variables: 74 for the gains, offsets, and transition parameters in the gait controller, and 16 for the parameters that set the model’s initial state. We set the CMA-ES parameters such that the population size of each generation (*λ*) was 16 and the update step for each generation used only the 8 best solutions (*μ*). Because these types of problems tend to have noisy, non-convex search spaces, and CMA-ES is a stochastic algorithm, we ran a set of multiple parallel optimizations with the same initial guess, and used the best solution from the set to seed the next set of optimizations as has been done previously [23,53]. Refer to S2 Table for the initial standard deviations used for each parameter. We chose to stop when the best solution from a set had an objective function value that was no better than 5% of the one from the previous set, as we observed optimized solutions that were within 5% of each other were qualitatively similar.

### Generating simulations of gait

To generate simulations of gait with a prescribed speed, we used 10 second simulations (*t*_des_ = 10 s) and a set of 20 parallel optimizations, each with a maximum of 3000 generations for CMA-ES. There were 7 simulations that were generated at prescribed speeds between 0.50 m/s and 2.00 m/s at an interval of 0.25 m/s. We first trained the model at the middle speed, 1.25 m/s, and then used those results to seed the neighboring speeds, continuing until we reached the lowest and highest speeds. For example, the solution for our 1.25 m/s case was used to seed the 1.00 m/s case, which was then used to seed the 0.75 m/s case.

To generate simulations of gait with a self-selected speed, we used longer, 30 second simulations (*t*_des_ = 30 s), as we found that solutions with a shorter desired simulation time would systematically fall forward at the end without a tight bound on step speed. We used a set of 10 parallel optimizations, each with a maximum of 1500 generations for CMA-ES. We generated 3 simulations of unimpaired gait at a self-selected speed, seeding from solutions of the slowest (0.50 m/s), middle (1.25 m/s), and fastest (2.00 m/s) cases from above. We also generated 18 simulations of impaired gait, starting with the unimpaired solution and seeding the next more severe case. For example, the solution from the unimpaired case was used to seed the mild SOL weakness case, which was used to seed the moderate case, which was finally used to seed the severe case.

## Acknowledgments

We would like to thank Apoorva Rajagopal and Jennifer Yong for their insightful feedback on this manuscript, and Christopher Dembia, Shrinidhi Lakshmikanth, Ajay Seth, and Thomas Uchida for helpful technical discussions. We thank the Stanford Research Computing Center for providing computational resources and support that contributed to these research results.

## Supporting information

**S1A Fig. Simulated walking at 0.55 m/s vs experimental data at 0.50 ± 0.20 m/s.** To facilitate comparison, normalized speed from experimental data was converted using the model’s leg length (0.856 m).

**S1B Fig. Simulated walking at 0.79 m/s vs experimental data at 0.84 ± 0.16 m/s.** To facilitate comparison, normalized speed from experimental data was converted using the model’s leg length (0.856 m).

**S1C Fig. Simulated walking at 1.04 m/s vs experimental data at 0.84 ± 0.16 m/s.** To facilitate comparison, normalized speed from experimental data was converted using the model’s leg length (0.856 m).

**S1D Fig. Simulated walking at 1.22 m/s vs experimental data at 1.24 ± 0.15 m/s.** To facilitate comparison, normalized speed from experimental data was converted using the model’s leg length (0.856 m).

**S1E Fig. Simulated walking at 1.46 m/s vs experimental data at 1.62 ± 0.18 m/s.** To facilitate comparison, normalized speed from experimental data was converted using the model’s leg length (0.856 m).

**S1F Fig. Simulated walking at 1.71 m/s vs experimental data at 1.62 ± 0.18 m/s.** To facilitate comparison, normalized speed from experimental data was converted using the model’s leg length (0.856 m).

**S1G Fig. Simulated walking at 1.96 m/s vs experimental data at 2.01 ± 0.27 m/s.** To facilitate comparison, normalized speed from experimental data was converted using the model’s leg length (0.856 m).

**S2 Fig. Key kinematic and kinetic parameters for all deficits.** This plot accompanies Fig 5 and shows key parameters for walking with simulated weakness (blue) and contracture (orange) of the SOL, GAS, or both (PF) along with simulated healthy gait (dotted line). Increasing color intensity indicates increasing deficit severity.

**S3A Fig. Simulated walking with mild, moderate, and severe soleus (SOL) weakness.**

**S3B Fig. Simulated walking with mild, moderate, and severe gastrocnemius (GAS) weakness. S3C Fig. Simulated walking with mild, moderate, and severe soleus and gastrocnemius (PF) weakness.**

**S3D Fig. Simulated walking with mild, moderate, and severe soleus (SOL) contracture.**

**S3E Fig. Simulated walking with mild, moderate, and severe gastrocnemius (GAS) contracture.**

**S3F Fig. Simulated walking with mild, moderate, and severe soleus and gastrocnemius (PF) contracture.**

**S1 Table. Muscle parameters for the unimpaired musculoskeletal model.**

**S2 Table. Initial Covariance Matrix Adaptation Evolutionary Strategy (CMA-ES) standard deviation for each free parameter.**

**S1 Video. Highlights.**

**S2 Video. Compilation of all simulations.**

