## Supplemental Figure 1 for "Predicting gait adaptations due to ankle plantarflexor muscle weakness and contracture using physics-based musculoskeletal simulations"

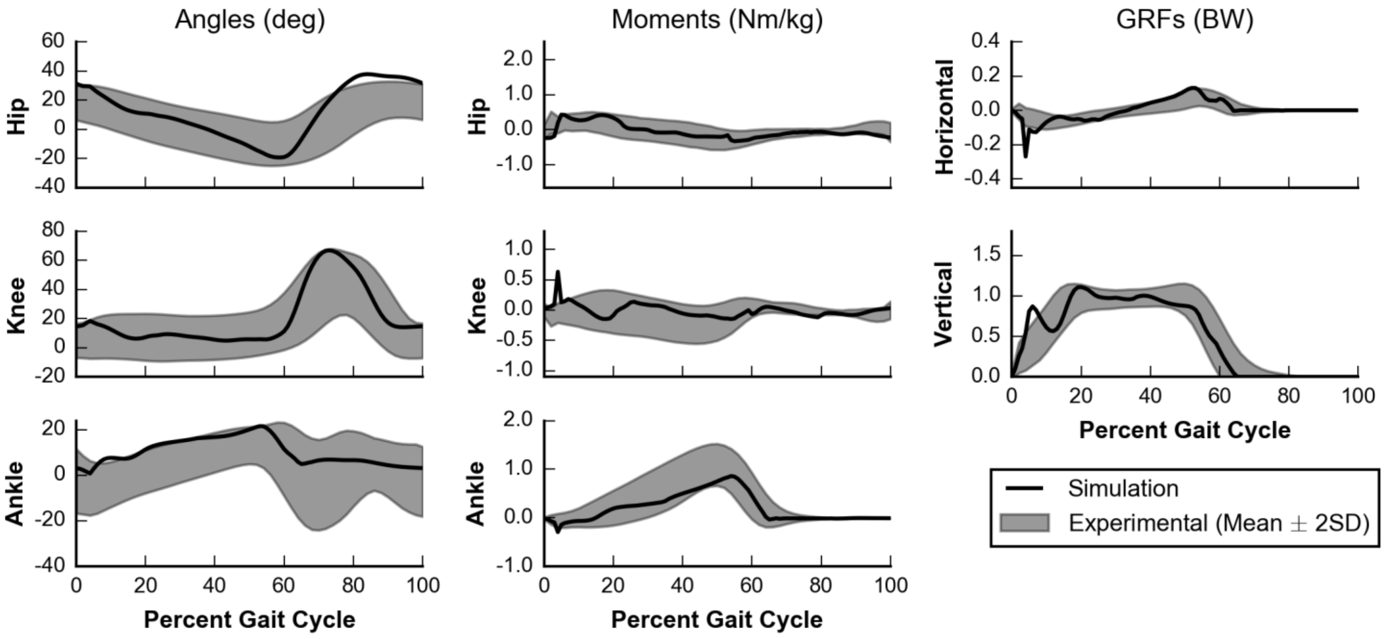


**S1A Fig. Simulated walking at 0.55 m/s vs experimental data at 0.50 ± 0.20 m/s.** To facilitate comparison, normalized speed from experimental data was converted using the model's leg length (0.856 m).

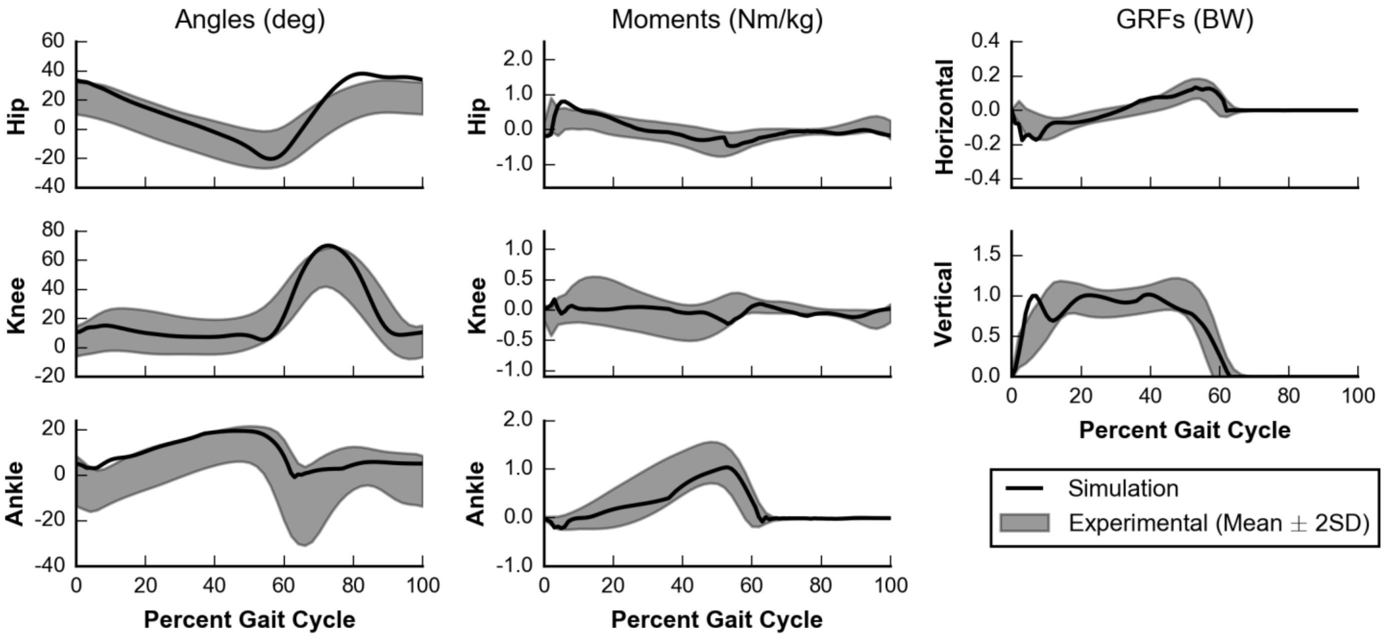


**S1B Fig. Simulated walking at 0.79 m/s vs experimental data at 0.84 ± 0.16 m/s.** To facilitate comparison, normalized speed from experimental data was converted using the model's leg length (0.856 m).

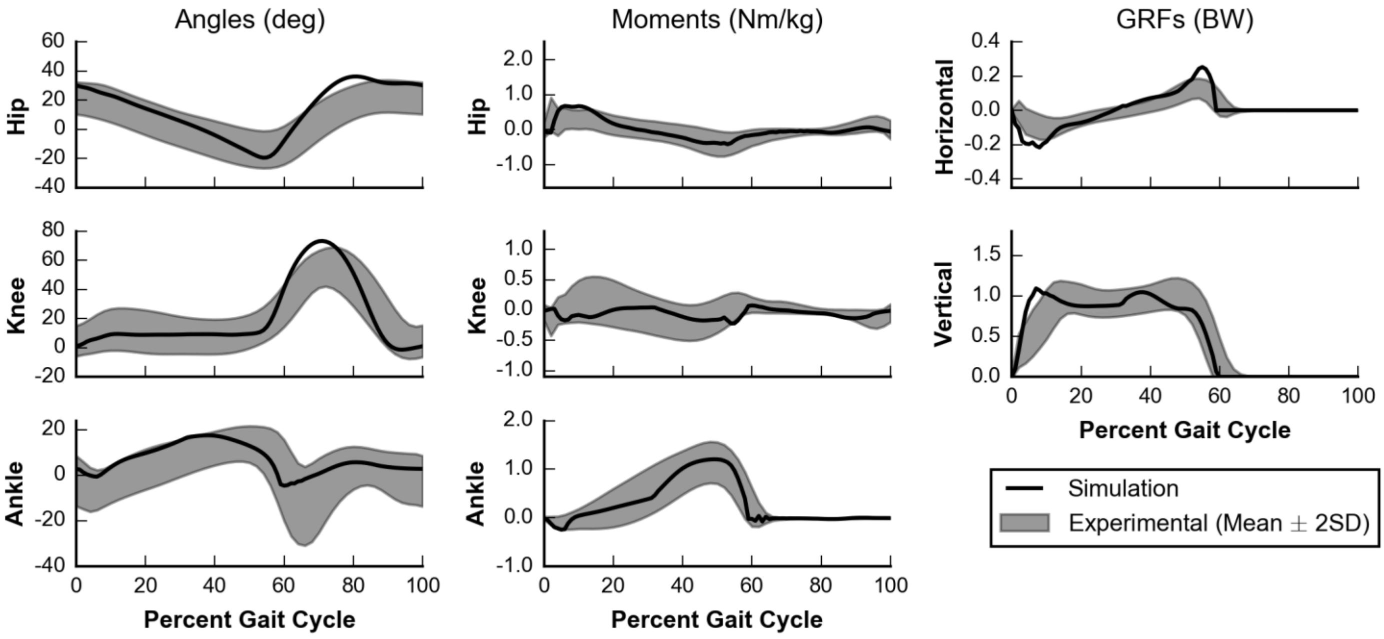


**S1C Fig. Simulated walking at 1.04 m/s vs experimental data at 0.84 ± 0.16 m/s.** To facilitate comparison, normalized speed from experimental data was converted using the model's leg length (0.856 m).

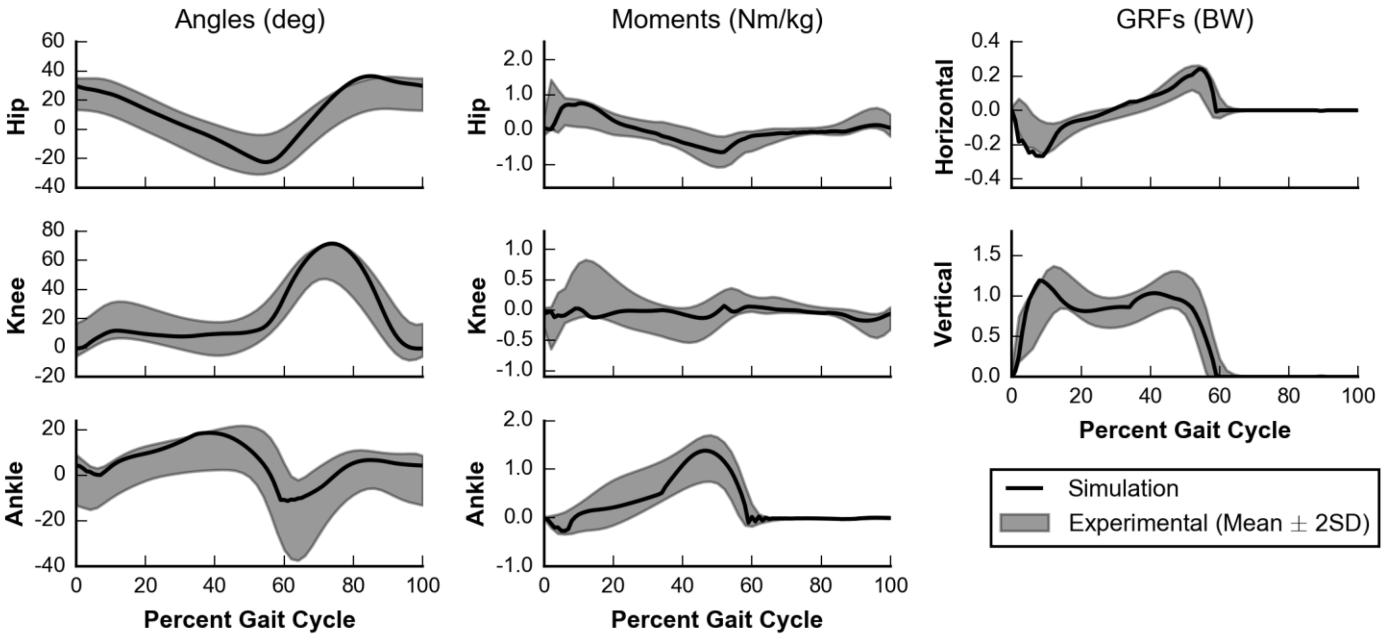


**S1D Fig. Simulated walking at 1.22 m/s vs experimental data at 1.24 ± 0.15 m/s.** To facilitate comparison, normalized speed from experimental data was converted using the model's leg length (0.856 m).

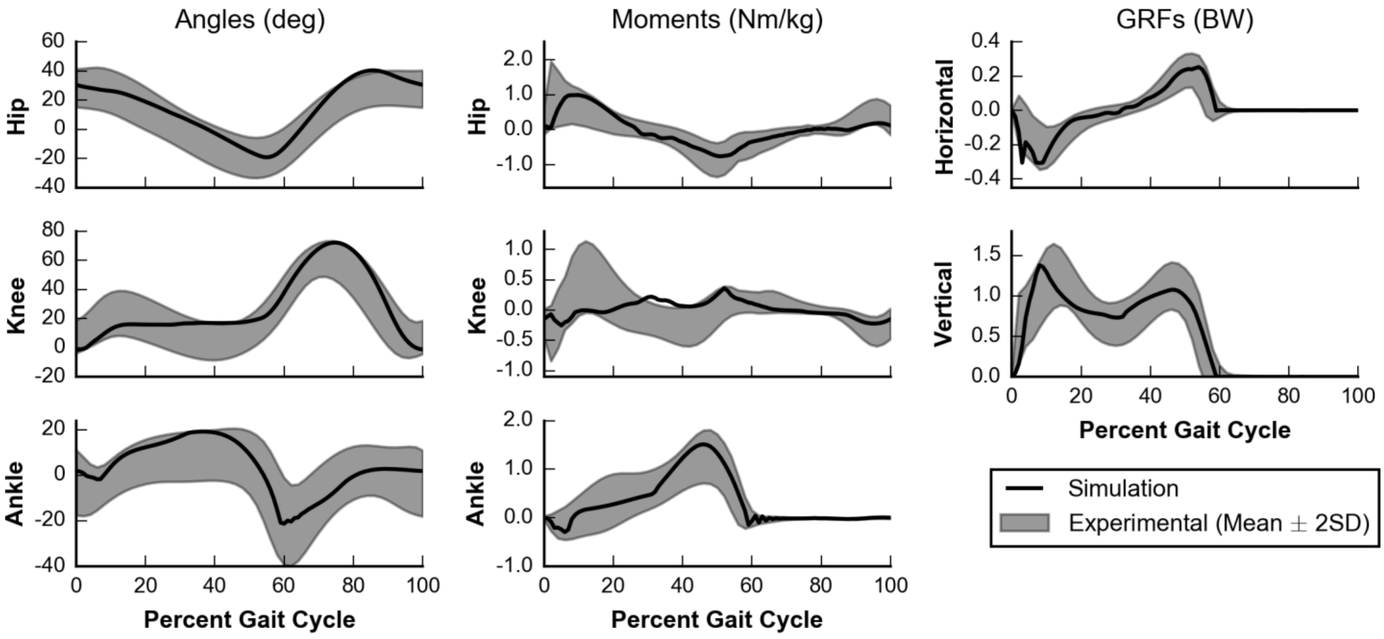


**S1E Fig. Simulated walking at 1.46 m/s vs experimental data at 1.62 ± 0.18 m/s.** To facilitate comparison, normalized speed from experimental data was converted using the model's leg length (0.856 m).

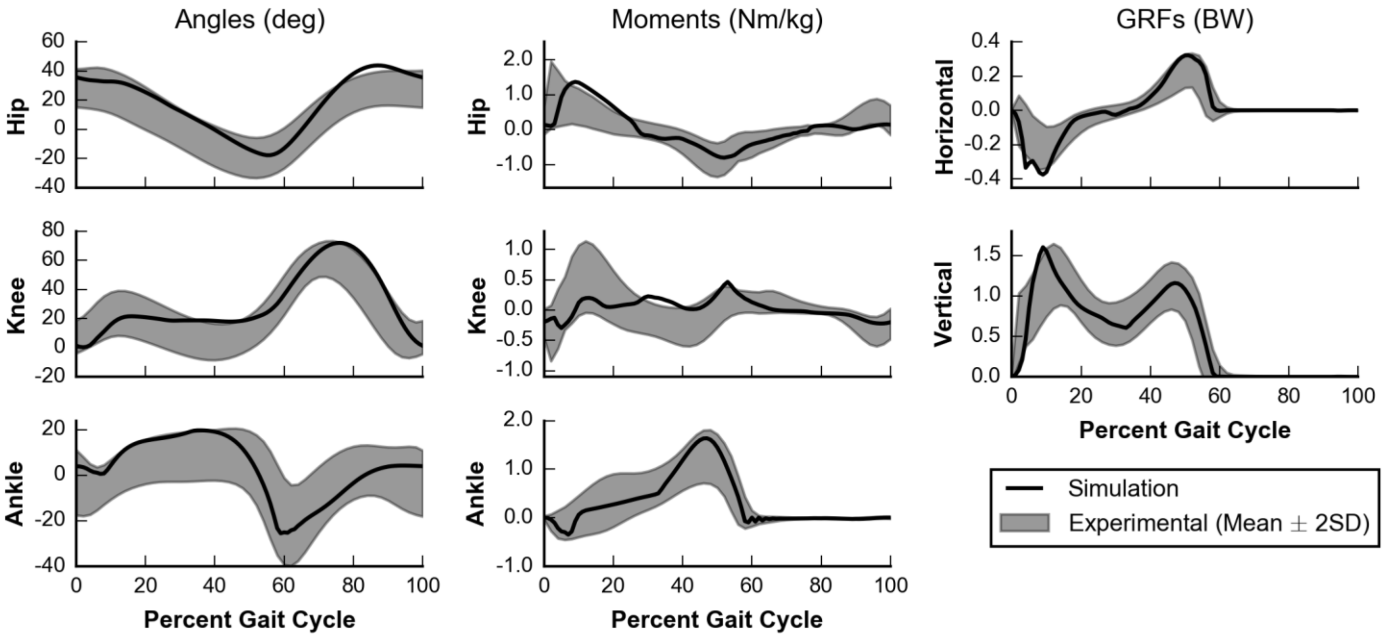


**S1F Fig. Simulated walking at 1.71 m/s vs experimental data at 1.62 ± 0.18 m/s.** To facilitate comparison, normalized speed from experimental data was converted using the model's leg length (0.856 m).

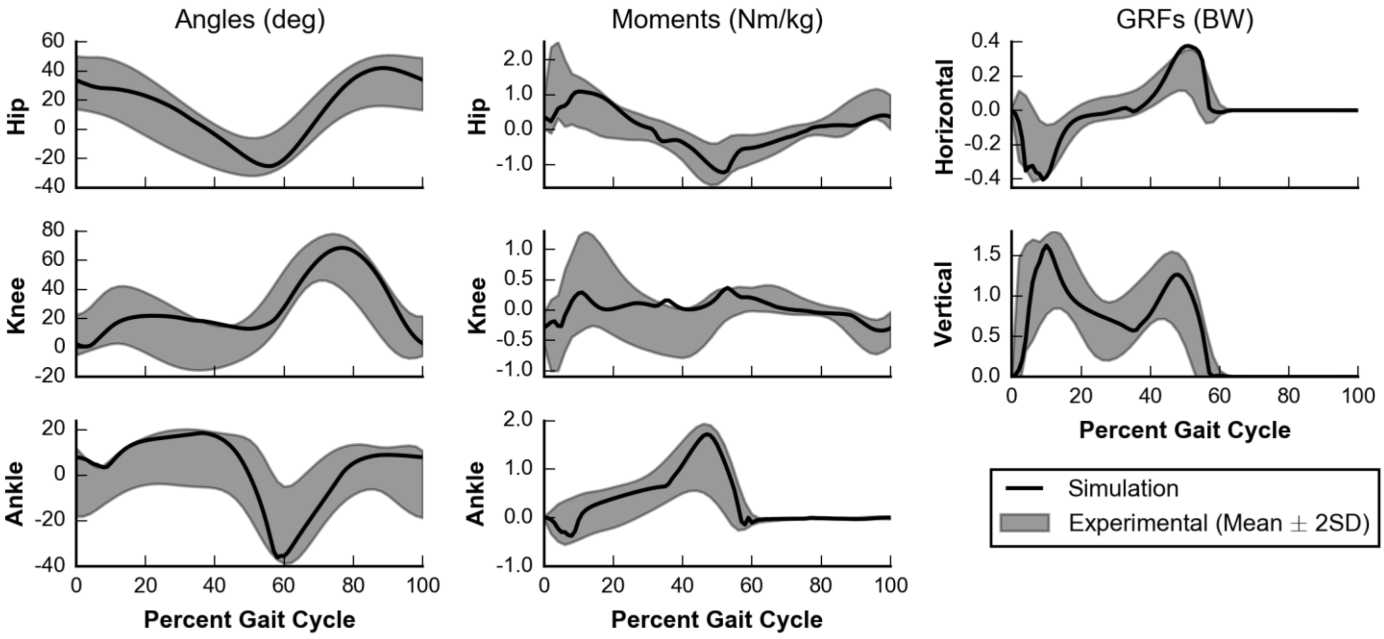


**S1G Fig. Simulated walking at 1.96 m/s vs experimental data at 2.01 ± 0.27 m/s.** To facilitate comparison, normalized speed from experimental data was converted using the model's leg length (0.856 m).
