## Supplemental Figure 2 for "Predicting gait adaptations due to ankle plantarflexor muscle weakness and contracture using physics-based musculoskeletal simulations"

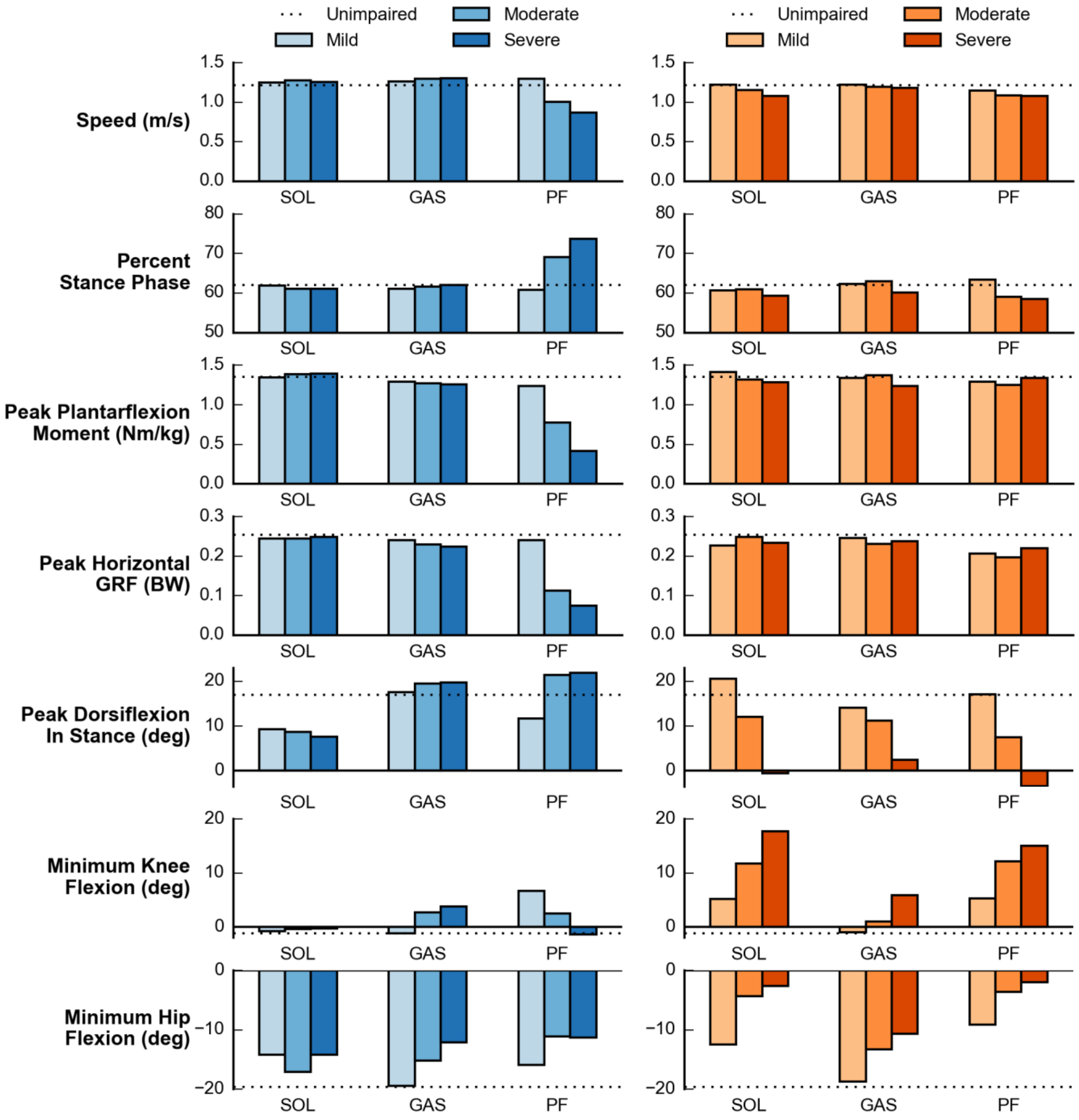


**S2 Fig. Key kinematic and kinetic parameters for all deficits.** This plot accompanies Fig 5 and shows key parameters for walking with simulated weakness (blue) and contracture (orange) of the SOL, GAS, or both (PF) along with simulated healthy gait (dotted line). Increasing color intensity indicates increasing deficit severity.
