## Supplemental Figure 3 for "Predicting gait adaptations due to ankle plantarflexor muscle weakness and contracture using physics-based musculoskeletal simulations"

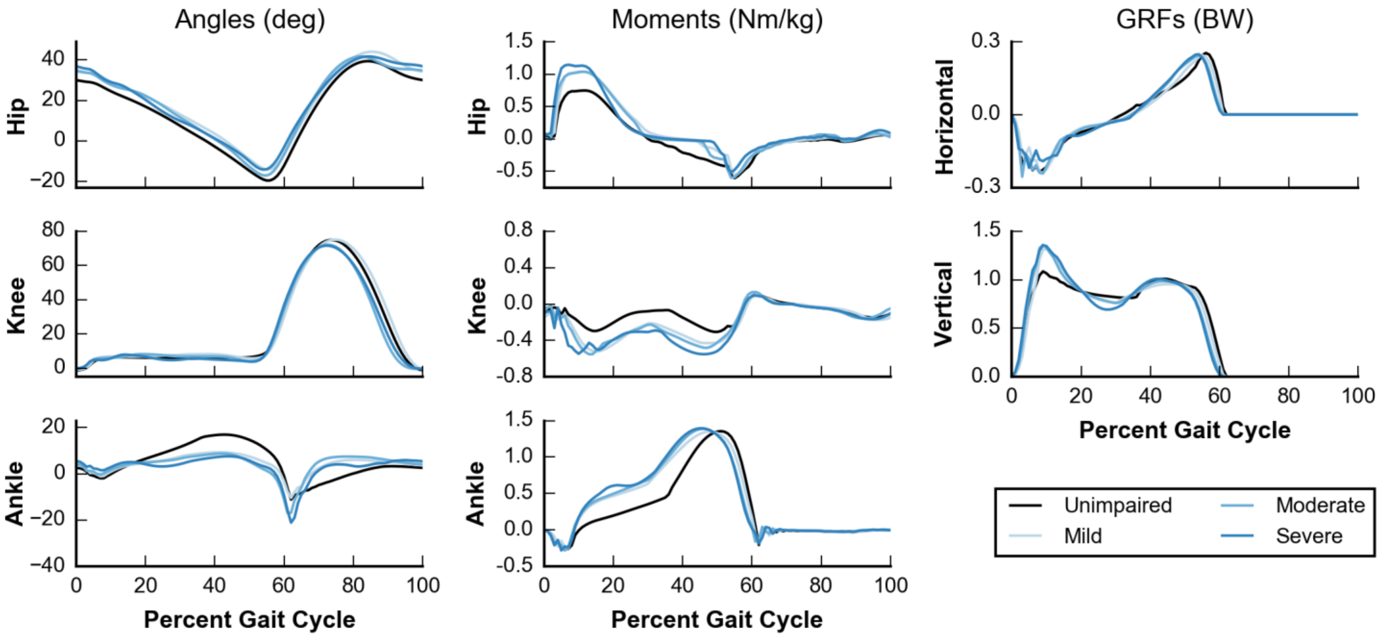


**S3A Fig. Simulated walking with mild, moderate, and severe soleus (SOL) weakness.**

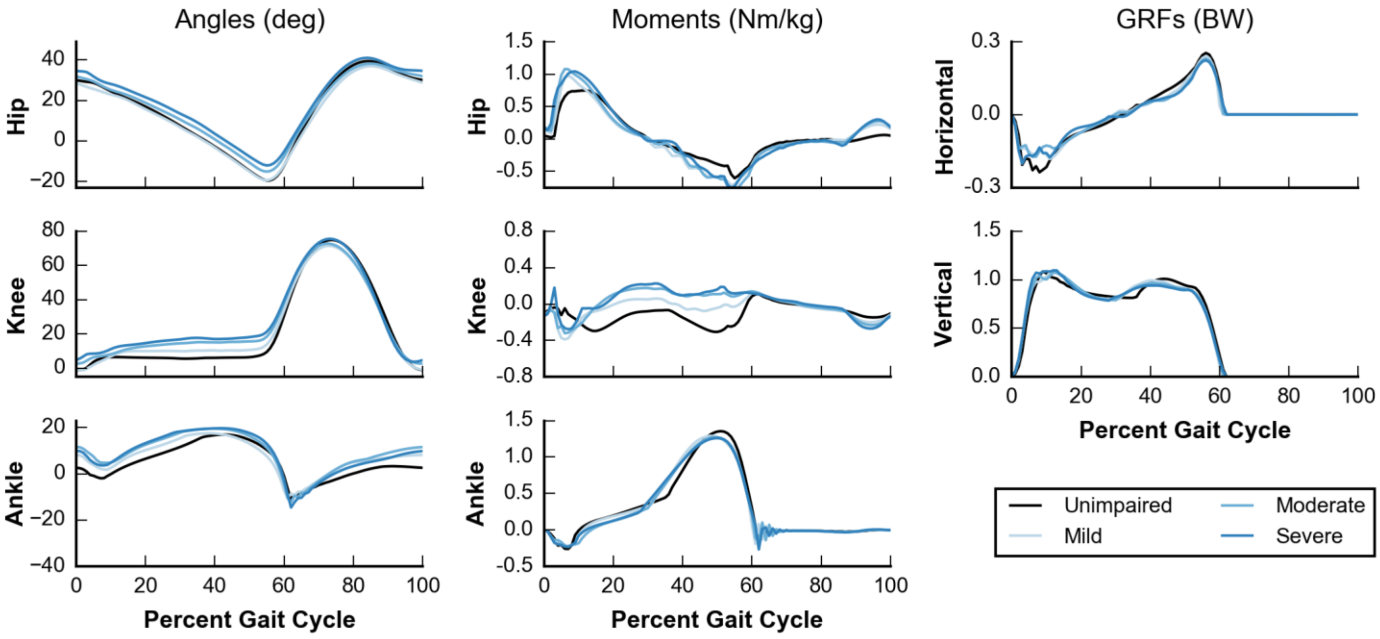


**S3B Fig. Simulated walking with mild, moderate, and severe gastrocnemius (GAS) weakness.**

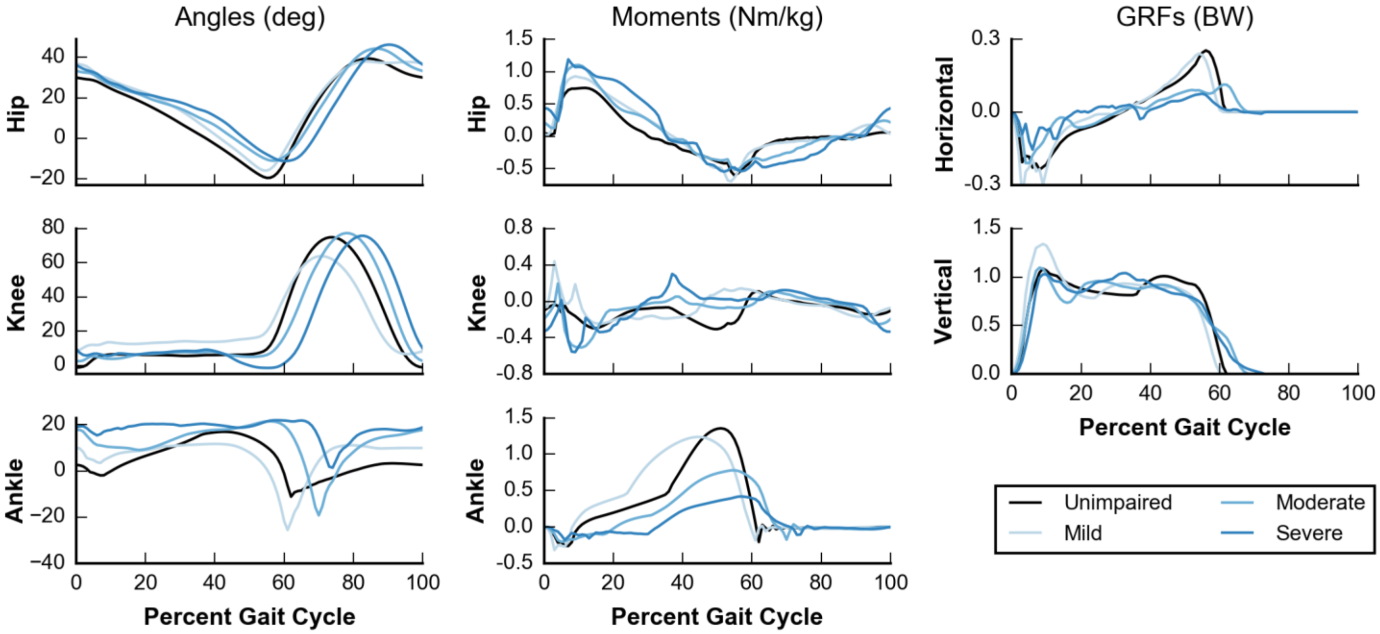


**S3C Fig. Simulated walking with mild, moderate, and severe soleus and gastrocnemius (PF) weakness.**

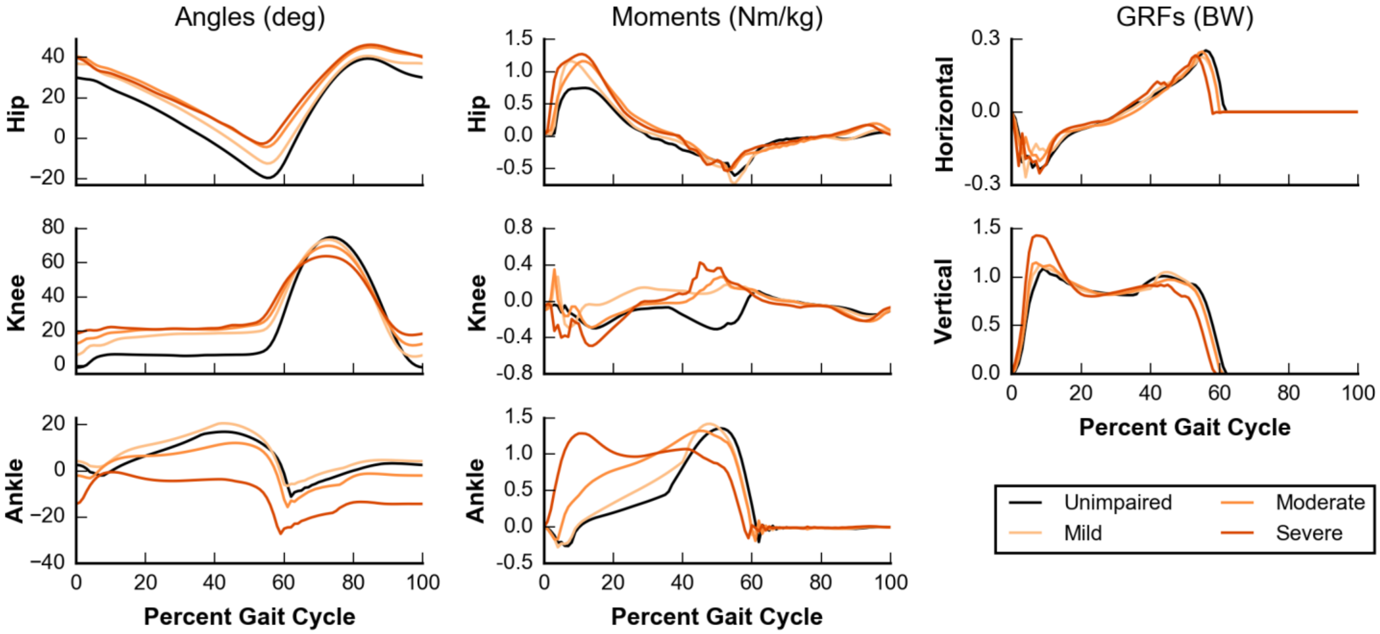


**S3D Fig. Simulated walking with mild, moderate, and severe soleus (SOL) contracture.**

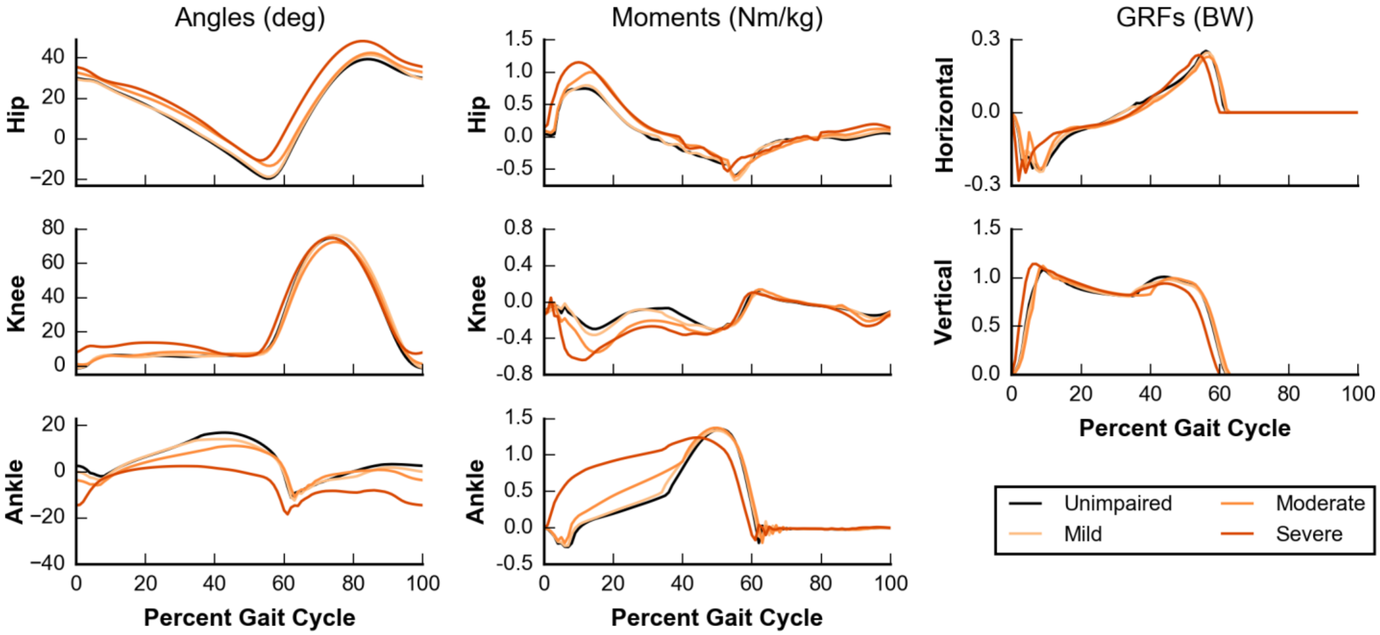


**S3E Fig. Simulated walking with mild, moderate, and severe gastrocnemius (GAS) contracture.**

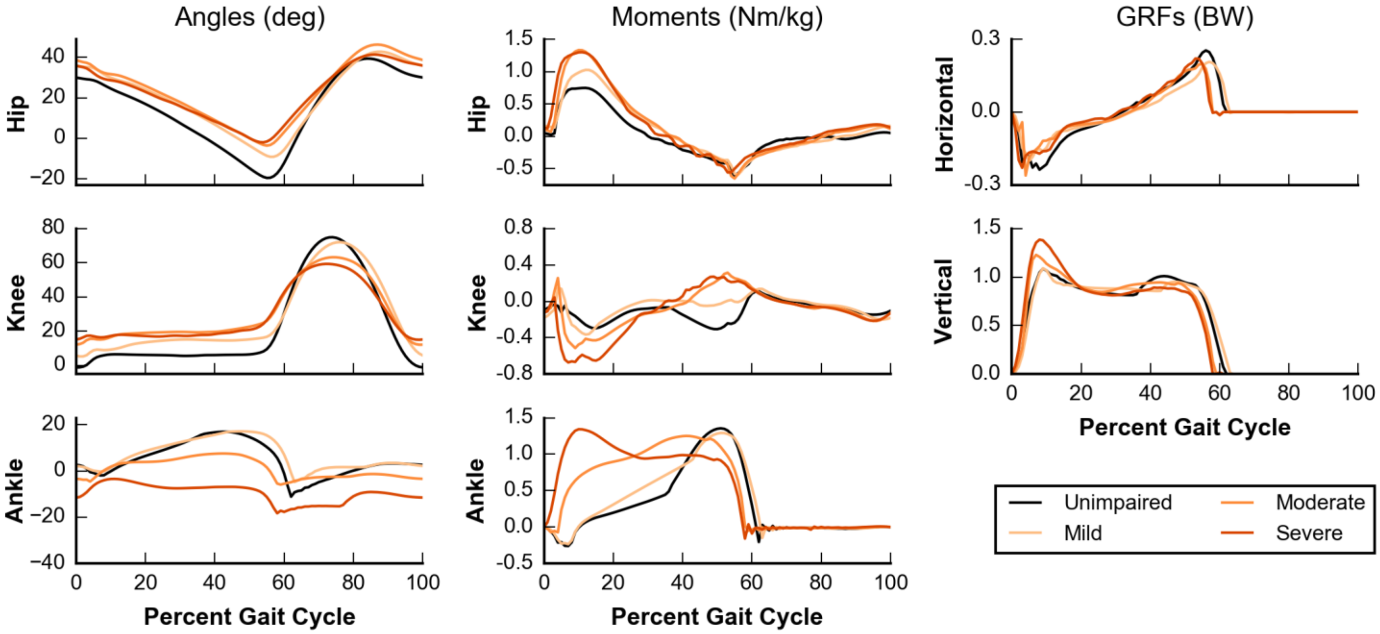


**S3F Fig. Simulated walking with mild, moderate, and severe soleus and gastrocnemius (PF) contracture.**
