## Supplemental Table 1 for "Predicting gait adaptations due to ankle plantarflexor muscle weakness and contracture using physics-based musculoskeletal simulations"

**S1 Table. Muscle parameters for the unimpaired musculoskeletal model.**

| Muscle name | Maximum isometric force ($F_o^m$)* [N] | Optimal fiber length ($l_o^m$) [m] | Muscle passive parameters ($k^{PE}$, $\varepsilon_0^m$)** | Maximum fiber contraction velocity [$l_o^m/s$] | Tendon slack length [m] | Tendon strain at $F_o^m$ | Muscle path from *Delp et al. 1990* model |
|---|---|---|---|---|---|---|---|
| **ILPSO** | 2697 | 0.117 | (5, 0.6) | 15 | 0.130 | 0.049 | psoas |
| **GMAX** | 3338 | 0.157 | (5, 0.6) | 15 | 0.048 | 0.049 | gluteus maximus2 |
| **RF** | 2192 | 0.076 | (9, 1.0) | 15 | 0.346 | 0.049 | rectus femoris |
| **HAMS** | 4105 | 0.069 | (5, 0.8) | 15 | 0.349 | 0.049 | semimembranosus |
| **VAS** | 9594 | 0.099 | (9, 1.0) | 15 | 0.102 | 0.049 | vastus intermedius |
| **BFSH** | 557 | 0.11 | (5, 0.6) | 15 | 0.117 | 0.049 | biceps femoris short head |
| **GAS** | 4691 | 0.051 | (5, 0.6) | 15 | 0.384 | 0.1 | medial gastrocnemius |
| **TA** | 2117 | 0.068 | (5, 0.6) | 15 | 0.238 | 0.049 | tibialis anterior |
| **SOL** | 7925 | 0.044 | (5, 0.6) | 15 | 0.244 | 0.1 | soleus |

* Maximum isometric force is based on a specific tension of 60 $N/cm^2$.

** $k^{PE}$ is the exponential shape factor for the passive force-length curve. $\varepsilon_0^m$ is the passive muscle strain at $F_o^m$. The symbols match those from *Thelen 2003*.
