## Supplemental Table 2 for "Predicting gait adaptations due to ankle plantarflexor muscle weakness and contracture using physics-based musculoskeletal simulations"

**S2 Table. Initial Covariance Matrix Adaptation Evolutionary Strategy (CMA-ES) standard deviation for each free parameter.**

| Parameter | Parameter description* | Initial CMA standard deviation |
|---|---|---|
| $K_C$ | constant excitation (Eq 1) | 0.01 |
| $K_{L+}$ | length feedback gain (Eq 2) | 0.1 |
| $l_o$ | length feedback offset (Eq 2) | 0.05 |
| $K_{V+}$ | velocity feedback gain (Eq 3) | 0.05 |
| $K_{F\pm}$ | force feedback gain (Eq 4) | 0.1 |
| $K_p$ | pelvis tilt orientation feedback gain (Eq 5) | 0.05 |
| $\theta_o$ | pelvis tilt orientation offset (Eq 5) | 0.01 |
| $K_v$ | pelvis tilt velocity feedback gain (Eq 5) | 0.05 |
| State transition: ES to MS | horizontal distance between ipsilateral foot and pelvis | 0.01 |
| State transition: PS to S | GRF on ipsilateral foot | 0.01 |
| State transition: S to LP | horizontal distance between ipsilateral foot and pelvis | 0.01 |
| State transition: LP to ES | GRF on ipsilateral foot | 0.01 |
| Initial positions | Initial pelvis tilt, hip, knee, and ankle angles | 0.01 |
| Initial velocities (except for pelvis horizontal velocity) | Initial pelvis tilt, hip, knee, and ankle angular velocities. Initial pelvis vertical translational velocity. | 0.01 |
| Initial pelvis horizontal velocity (targeted speed)** | Initial pelvis horizontal velocity during a simulation with a targeted speed. | 0.01 |
| Initial pelvis horizontal velocity (self-selected speed)** | Initial pelvis horizontal velocity during a self-selected speed simulation. | 0.1 |

* Equation numbers in this column refer to those in the main text.

** Since these two affect the same parameter, only one of these is used depending on the type of simulation (i.e., targeted speed or self-selected speed).
